# Metabolically-Driven Maturation of hiPSC-Cell Derived Cardiac Chip

**DOI:** 10.1101/485169

**Authors:** Nathaniel Huebsch, Berenice Charrez, Brian Siemons, Steven C. Boggess, Samuel Wall, Gabriel Neiman, Verena Charwat, Karoline H. Jæger, David Cleres, Åshild Telle, Felipe T. Lee-Montiel, Nicholas C. Jeffreys, Nikhil Deveshwar, Andrew Edwards, Jonathan Serrano, Matija Snuderl, Andreas Stahl, Aslak Tveito, Evan W. Miller, Kevin E. Healy

## Abstract

Human induced pluripotent stem cell derived cardiomyocytes (hiPSC-CM) are a promising *in vitro* tool for drug development and disease modeling, but their immature electrophysiology limits diagnostic utility. Tissue engineering approaches involving aligned 3D cultures enhance hiPSC-CM structural maturation but are insufficient to induce mature electrophysiology. We hypothesized that mimicking post-natal switching of the heart’s primary ATP source from glycolysis to fatty acid oxidation could enhance electrophysiological maturation of hiPSC-CM. We combined hiPSC-CM with microfabricated culture chambers to form 3D cardiac microphysiological systems (MPS) that enhanced immediate microtissue alignment and tissue specific extracellular matrix (ECM) production. Using Robust Experimental design, we identified a maturation media that improved calcium handling in MPS derived from two genetically distinct hiPSC sources. Although calcium handling and metabolic maturation were improved in both genotypes, there was a divergent effect on action potential duration (APD): MPS that started with abnormally prolonged APD exhibited shorter APD in response to maturation media, whereas the same media prolonged the APD in MPS that started with aberrantly short APD. Importantly, the APD of both genotypes was brought near the range of 270-300ms observed in human left ventricular cardiomyocytes. Mathematical modeling explained these divergent phenotypes, and further predicted the response of matured MPS to drugs with known pro-arrhythmic effects. These results suggest that systematic combination of biophysical stimuli and metabolic cues can enhance the electrophysiological maturation of hiPSC-derived cardiomyocytes. However, they also reveal that maturation-inducing cues can have differential effects on electrophysiology depending on the baseline phenotype of hiPSC-CM. *In silico* models provide a valuable tool for predicting how changes in cellular maturation will manifest in drug responsiveness.

## Introduction

Human induced pluripotent stem cell (hiPSC) technology provides an exciting opportunity for disease modeling and drug discovery. An immediate goal for cardiac tissue models formed from hiPSC-derived cardiomyocytes (hiPSC-CM) is to reduce and refine the use of animal testing in the drug development pipeline^1^. Inherent differences between species have historically diminished the ability of animal models to prognosticate drug safety and efficacy. Microphysiological systems (MPS), or organ-chips, combine 3D-architecture of tissue micro-environments with the ability to interrogate key physiological functions (for example, cardiomyocyte action potential) and well-defined delivery profiles for nutrients.

A challenge with using hiPSC-CM to predict drug safety and efficacy is the immaturity of hiPSC-CM^2–5^. In particular, hiPSC-CM exhibit automaticity (spontaneous beating without electrical stimulation) and longer action potentials (445±73msec for ventricular-like hiPSC-CM, versus 270-300msec directly measured by patch-clamp of primary human adult left-ventricular cardiomyocytes)^6–11^. Culturing hiPSC-CM or human embryonic stem cell (hESC) derived cardiomyocytes (hESC-CM) within the *in vivo*-like micro-environment of Engineered Heart Muscle (EHM^12–21^), or MPS^22^ has been shown to mature hiPSC-CM to some extent by enhancing physiologic hypertrophy, and lead to pharmacology more closely correlated to the one of the adult human heart. However, these approaches have not led to consistent electrophysiological maturation of hiPSC-CM.

In addition to 3D culture approaches, bioreactor-based strategies such as chronic electrical pacing or cyclic strain (typically applied over 2-4 weeks in culture) enhance maturity of hESC-CM and hiPSC-CM^15,18,23,24^. Collectively these methods are promising, but no single approach applied thus far has been sufficient to induce full maturation of pluripotent stem cell derived cardiomyocytes. Furthermore, many of the existing approaches require prolonged culture periods (in some cases approaching one month), large scale formats, and/or complex setups to execute. These issues would lead to cost and logistical limitations for their translation to higher-throughput analyses that would be essential to use these technologies for drug development.

As bioreactor approaches have limited scalability, and tissue engineered micro-environments alone are not sufficient to induce hiPSC-CM maturation consistent with the adult heart, there has been a focus on combining tissue engineering approaches with soluble cues. Consistent with the notion that micro-environmental cues can enhance hiPSC-CM maturity, transplanting hiPSC-CM into neonatal rodent hearts enhances maturation^25^. One possible explanation for this result is that the milieu of the heart contains soluble cues that enhance cardiomyocyte maturity. Reductionist approaches have used specific soluble cues from the fetal and post-natal heart, including cytokines^26,27^, micro-RNAs^28^, heart specific extracellular matrix^29^ and hormones^30^ to enhance the maturity of hESC-CM and hiPSC-CM.

Metabolic cues like glucose levels provide another key facet of the heart’s soluble environment that may be particularly important to cardiomyocyte maturation. Postnatally, the heart switches from glycolysis to fatty-acid oxidation as its primary source of ATP^31,32^. Previously, 2D hiPSC-CM monolayers and engineered tissues exposed to glucose-depleted, fatty-acid enriched media exhibited more mature metabolic profiles and physiology compared to hiPSC-CM cultured in standard media^33–36^. However, fatty-acid based media has not been studied in the context of MPS, and potential population variance in cellular response to fatty-acid media has not been explored. We hypothesized that the combination of aligned, 3D culture and fatty-acid could enhance electrophysiological maturation of hiPSC-CM within cardiac MPS. We investigated hiPSC-CM derived from two distinct donor lines labeled WTC^37^ and SCVI20^38^. Both lines were derived from healthy individuals who did not harbor known cardiomyopathy-associated mutations. Using a Design of Experiments approach^39–41^, in MPS derived from WTC hiPSC, we identified fatty-acid based Maturation Media (MM) that led to a more mature metabolic phenotype and improved calcium handling in MPS. These changes were associated with shortened, adult-like Action Potential Duration (APD) in WTC-MPS. MM-treated MPS also exhibited changes in expression of ion-channel and calcium handling genes, including Sarcolipin (SLN).

In contrast, MM had different effects on MPS derived from SCVI20 hiPSC. At baseline, monolayers and SCVI20 hiPSC-CM - derived MPS had extremely short action potentials. Whereas MM markedly shortened APD in WTC MPS, the same media prolonged APD in SCVI20 MPS. In both cases, MM treatment led to APD within the range of 270-300msec expected for adult human left-ventricular cardiomyocytes. In both genotypes, MM produced metabolic maturation, and a trend toward a more robust calcium transient (ΔF/F_0_) and enhanced contractility at low extracellular calcium. To understand these differential responses to common media, we used mathematical modeling to predict the contribution of specific current fluxes and calcium handling machinery to action potentials and calcium transients. These models predicted that individual currents were very different between immature MPS of different genotypes, despite both basal hiPSC having been derived from healthy male volunteers who did not harbor genetic diseases. MM treatment led to a convergent set of ion currents and calcium handling parameters, and the MM-induced changes in simulated currents were consistent with shifts in the pharmacology of MM versus Standard Media (SM). These results suggest that maturation media can both improve overall tissue physiology and minimize artificial differences in pharmacology of hiPSC-CM due to non-genetic sources of variability (e.g. clonal variation). Potentially, this can subsequently reveal true pharmacogenomic relationships.

## Results and Discussion

### Robust Design Experiments Indicate Optimal Carbon Sourcing for Mature Beating Physiology in WTC-Derived hiPSC-Cardiac MPS

We optimized fabrication of hiPSC-CM derived cardiac MPS^22^ to enhance the consistency of multilayer tissue self-assembly (**Fig. 1A-C**) and subsequent achievement of aligned sarcomeres and uniaxial beating **(Fig. 1D,E)**. To minimize batch-to-batch variability in cellular composition that might hinder drug screening studies, we formed cardiac MPS that mimicked the mass composition of the human heart by combining a defined cocktail^42^ of 80% WTC hiPSC-CM and 20% isogenic hiPSC-SC (**Supplemental Methods, Fig. S1,S2**). We employed Robust Experimental Design to screen for the effects of glucose, oleic acid, palmitic acid, and albumin (bovine serum albumin, BSA) levels on WTC MPS maturity (**Table 1**). MPS were incubated with different fatty-acid media for ten days before their beating physiology and calcium flux were assessed. Optimal media would reduce automaticity (e.g. reduce spontaneous beating rate), while reducing the interval between peak contraction and peak relaxation (a surrogate for APD) in field-paced tissues (**Fig. 2A-C**), and maintaining a high level of beating prevalence during pacing (defined as the percent of the tissue with substantial contractile motion; **Fig. 2D**). Beating interval correlated with rate-corrected Full-Width Half Maximum calcium flux time, FWHM (**Fig. 2B,E,G**), and beating prevalence correlated with calcium flux amplitude (**Fig. 2D,F,G**). The strong correlations between calcium flux and contractile motion with respect to timing and amplitude suggest that none of the medias used in this screen disrupt calcium-contraction coupling. However, a concurrent decrease in spontaneous beat rate and reduction in beating interval suggests a diminished leak current or lower resting membrane potential; both are concurrent with more mature electrophysiology.

**Figure 1.**
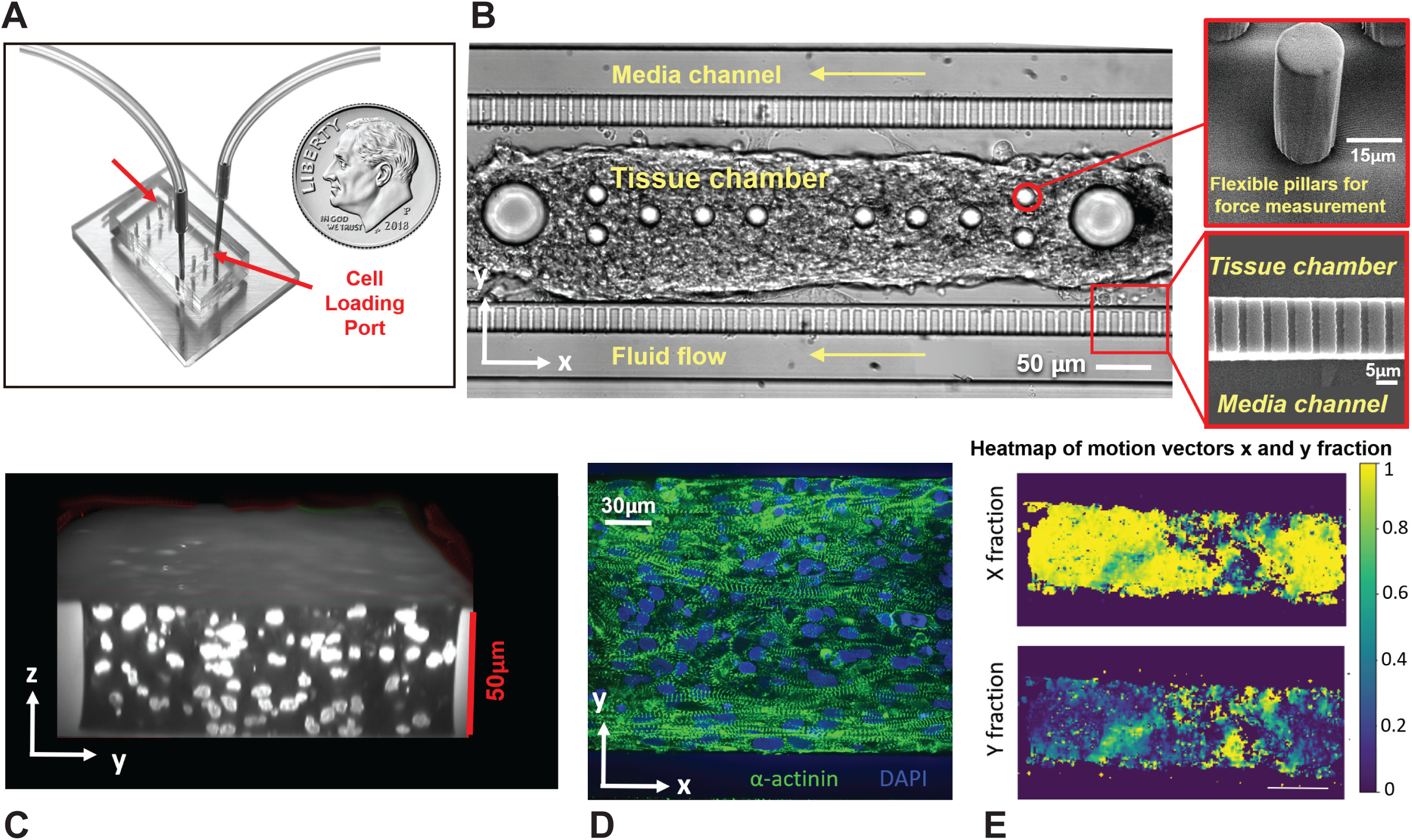
Optimized hiPSC-CM Microphysiological Systems (MPS). **A)** Representative image of two cardiac chips parallel to one another. The fluidic inlet and outlet, coupled via tubing, flank the cell loading ports. **B)** Representative brightfield image of a cardiac chip, showing cell loading chamber surrounded by media channels, with accompanying SEM images of flexible pillars for *in situ* contraction force measurements (inset top), and fenestrations insuring diffusive transport of nutrients (inset bottom). **C)** Representative confocal micrograph depicting several layers of cell thickness (side view of DRAQ5 stained nuclei). **D)** Representative confocal micrograph of a cardiac MPS indicating robust sarcomere alignment (sarcomeric α-Actinin, green) and **C)** Heatmap of motion vectors obtained through motion tracking of 8×8 pixel macroblocks overtime. Both length (X) and width (Y) direction of motion vectors are shown, indicating that 95% of the contraction coincides with the orientation of the X axis. These motion vectors were used to analyze the prevalence of beating (center; percent of the tissue that moves with average speed above a defined threshold that was held constant for all tissues).

**Table 1.** Design of Experiments (L9) for Media Screening.

| **Media Formulation** | **Glucose Concentration (mM)** | **Oleic Acid Concentration (μM)** | **Palmitic Acid Concentration (μM)** | **Bovine Serum Albumin Concentration (mg/mL)** |
| --- | --- | --- | --- | --- |
| **1\* (Standard Media, SM)** | 25 | 0 | 0 | 2.5<sup>++</sup> |
| **2** | 25 | 62.5 | 100 | 10 |
| **3** | 25 | 125 | 200 | 25 |
| **4** | 10 | 62.5 | 0 | 25 |
| **5** | 10 | 125 | 100 | 2.5 |
| **6** | 10 | 0 | 200 | 10 |
| **7<sup>&</sup>** | 0 | 125 | 0 | 10 |
| **8<sup>&</sup>** | 0 | 0 | 100 | 25 |
| **9<sup>&</sup>** | 0 | 62.5 | 200 | 2.5 |
| **10<sup>&</sup> (Maturation Media, MM)** | 2.75 | 125 | 100 | 25 |
| **M1<sup>&</sup>** | 2.75 | 125 | 0 | 2.5 |
| **M2<sup>&</sup>** | 2.75 | 125 | 0 | 25 |
| **M3<sup>&</sup>** | 2.75 | 125 | 100 | 2.5 |
| **M4<sup>&</sup>** | 2.75 | 125 | 100 | 10 |

\* The composition of Media 1 is identical to standard RPMI used to feed hiPSC-CM. All media were prepared from the same batch of powdered, glucose free RPMI.
++ The B-27 supplement contains 2.5mg/mL BSA; thus, in media formulations containing only 2.5mg/mL BSA, no extra albumin was added to the system.
& Medias without glucose, and MM, were supplemented with 10mM galactose
In addition to the ingredients described above, all media were supplemented with 2% B-27 supplement and 150μg/mL *L*-ascorbic acid.

**Figure 2.**
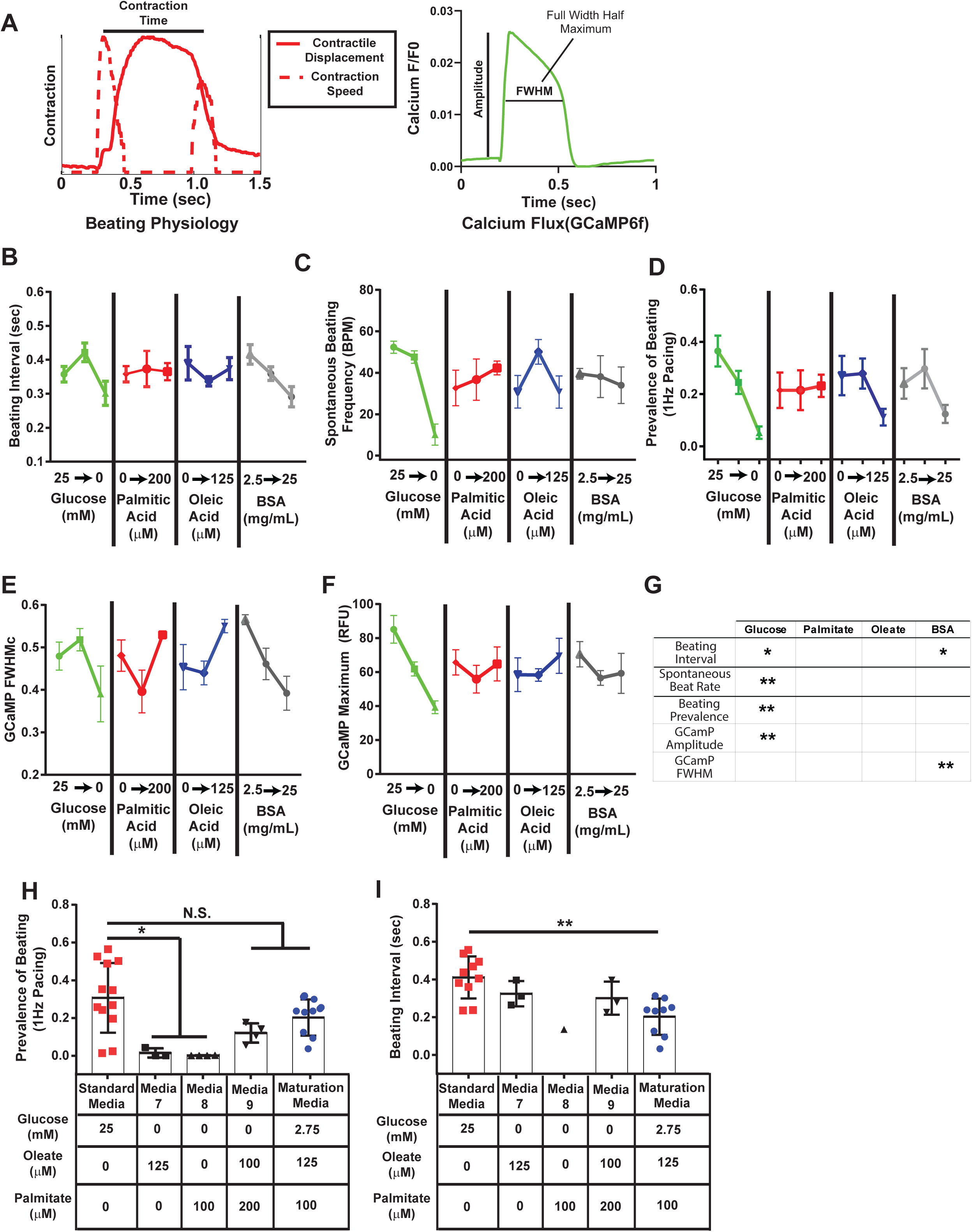
Design-of-Experiments (DoE) Based Screens Identify Maturation Media for hiPSC-CM Microphysiological Systems. **A)** Approach used for initial screen. Computational motion capture is performed on bright-field videos of contracting cardiac MPS, giving the contraction time (defined as the distance between peaks in motion speed for contraction and relaxation, which approximates the interval over which displacement occurs). The knock-in reporter, GCaMP6f, is used to monitor the timing (rate corrected Full-Width-Half-Maximum, FWHM and amplitude of calcium transients in MPS. **B-F)** Results from representative L9 Taguchi Array experiments, depicting **B)** beating interval, **C)** spontaneous beating frequency, and **D)** tissue beating prevalence, all obtained from motion tracking analysis, along with calcium transient **E)** FWHM and **F)** amplitude, obtained from analysis of GCaMP6f fluorescence. **G)** Summary L9 analysis. 1-way ANOVA tests were performed to assess the effects of specific media components. Increases in the parameters measured were denoted as * or ** for *p*-values of 0.05 and 0.01, respectively. **H-I)** Comparison of MPS cultured in standard media (red) to MPS cultured in glucose-free media during L9 studies (black), and MPS cultured in the final Maturation Media (blue). MPS were examined for the effects of glucose depletion and fatty acid addition on **H)** beating prevalence and I) beating interval. Note, beating prevalence and calcium amplitude are ideally maximized, while beating interval, spontaneous beat rate, and calcium transient FWHM, are ideally minimized, in adult left ventricular cardiomyocytes. MPS were cultured for ten days prior to analysis for the L9 experiments. Data: **B-F**: plot of mean ± SEM, *n* = 9; **H-I**: all data points with median, *n* = 3-12, except for beating interval in media 8, which could only be calculated in one sample (no other samples cultured in this media exhibited either spontaneous or paced beating) ** *p* < 0.01, * *p* < 0.05 (2-way *t*-test with Holm-Bonferonni correction for multiple comparisons).

Screening experiments did not suggest a significant role for either oleic or palmitic acid alone in inducing shortened beating intervals (**Fig. 2B**), and trends toward increased calcium transient duration at intermediate levels of the fatty acids were not statistically significant (**Fig. 2E**). However, these fatty acids did have effects on beating physiology in the context of concurrent glucose deprivation: completely omitting glucose while adding in galactose in MPS treated with media having only oleic acid or palmitic acid eliminated beating under 1Hz pacing. In contrast, treatment with glucose-free (galactose containing) media that was supplemented with both fatty acids (Media 9; **Fig. 2H**) partially rescued this deficiency. This is consistent with the ability of hiPSC-CM to use both these fatty acids as ATP sources^35^. Thus, we concluded that the optimal media should include both palmitic and oleic acids. Furthermore, although absolute glucose deprivation would likely force fatty acid oxidation, and previous work has established galactose combined with fatty acids as a viable ATP source for healthy hiPSC-CM in monolayer cultures^35,43^, we observed that in MPS, complete glucose deprivation dramatically reduced beating prevalence and calcium transient amplitude, even in the presence of 10mM galactose. This prompted us to adjust the glucose level media to a low but non-zero level of 0.5g/L (2.75mM; ∼10% of the level in standard RPMI Media).

As the inclusion of higher levels of BSA appeared to diminish beating interval without severely affecting prevalence or calcium flux, we concluded that an ideal maturation media would contain this higher level (2.5%, vs. 0.25% contained in standard, B-27 supplemented media; **Fig. 2B,E**). This led to a new Maturation Media (herein referred to as “MM”): glucose free RPMI basal media supplemented with 2% B-27, 0.5g/L glucose (2.8mM), 10mM galactose, 2.25% BSA (to a final concentration of 2.5% BSA, including the BSA contained in B-27), 125µM oleic acid and 100µM palmitic acid. MM exhibited a substantial portion of the beneficial effects of glucose free, fatty acid enriched media on reducing beating interval, without a concurrent loss of beating prevalence with 1 Hz pacing (**Fig. 2H, I**).

### Maturation-Media Induced Changes in Action Potential and Calcium Transients for WTC MPS

In WTC MPS, MM reduced APD from the prolonged levels we observed for Standard Media (RPMI containing B-27 supplement; SM) treated MPS (**Fig. 3A,B,E**). Interestingly, however, switching from SM to MM had no measurable effects on APD or automaticity when hiPSC-CM were cultured in confluent 2D monolayers (**Fig. 3E, S3E,F**). GCaMP6f based analysis of Ca^2+^ handling revealed that baseline-normalized calcium amplitude (ΔF/F_0_) was significantly increased by fatty-acid based MM in WTC MPS **(Fig. 3G)**. However, we did not observe a significant change in the maximum calcium upstroke velocity in paced tissues (**Fig. 3H**). Likewise, action potential upstroke timing was not significantly affected by MM pre-treatment, although there was a trend toward shorter action potential upstroke times in MM pre-treated WTC MPS and hiPSC-CM monolayers (**Fig. S3A**).

**Figure 3.**
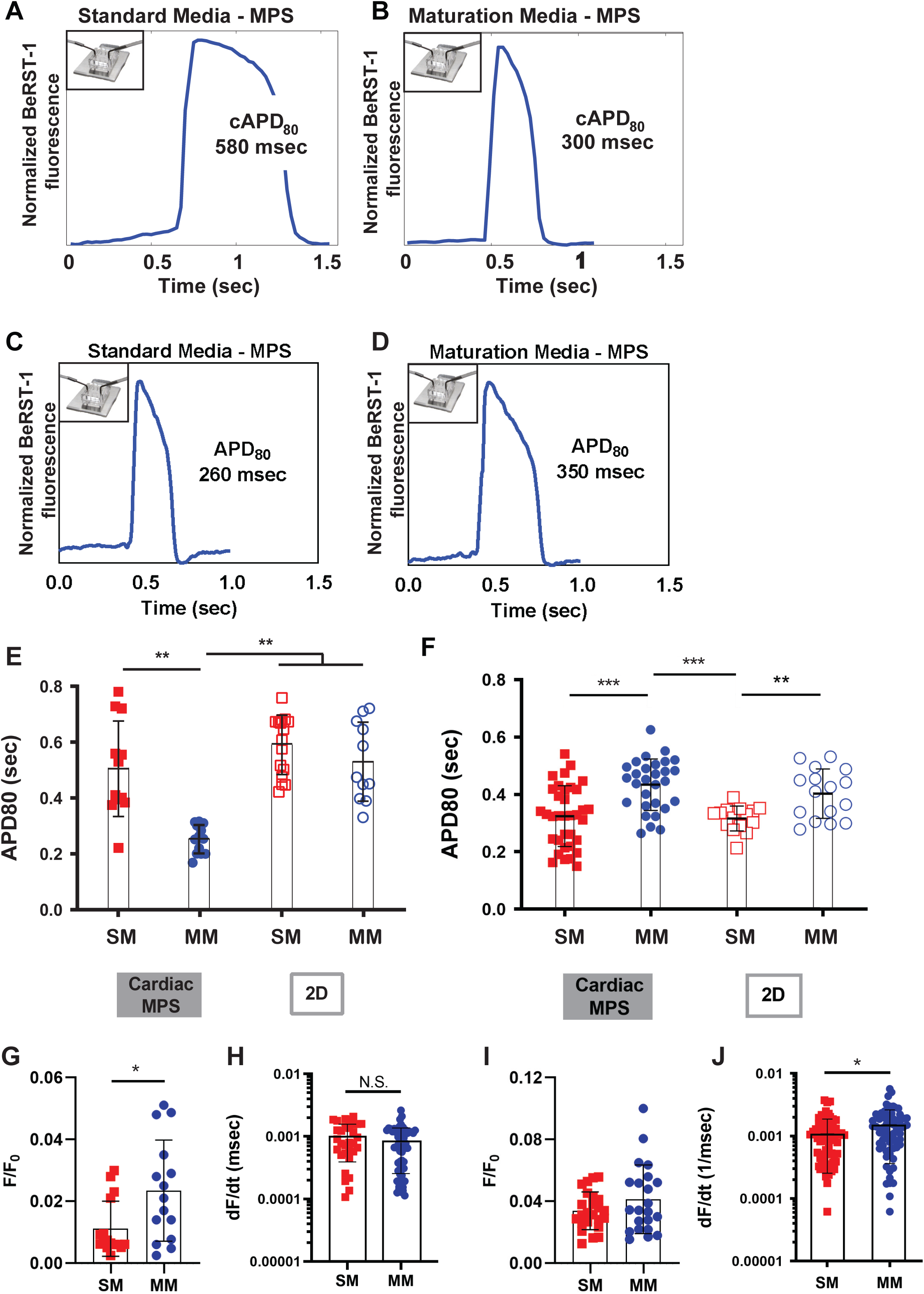
Action Potential Characterization of Matured Cardiac MPS. **A-D)** Representative voltage tracings for **(A,B)** WTC MPS and (**D,E)** SCVI20 MPS, cultured for ten days in either (**A,C**) standard cardiac media, or (**B,D**) Maturation Media (MM). Voltage tracings were obtained by overnight labeling of MPS with BeRST-1. **E,F)** Quantitative analysis of 80% action potential duration (APD_80_) for **E**) WTC or **F)** SCVI20 MPS (closed shapes) and monolayers (open shapes), cultured in standard cardiac media (SM; red) or maturation media (blue). **G-J)** Background normalized calcium amplitude of **G**) WTC and **I**) SCVI20 cell lines, and maximum Ca^2+^ upstroke velocity of **H)** WTC and **I**) SCVI20 lines. All data: plot of all points with median, *n* > 5. (* *p* < 0.05, ** *p* < 0.01, *** *p* < 10^−3^; 2-way *t*-test with Holm-Bonferonni correction for multiple comparisons).

Due to concerns about potential cytotoxicity of high levels of BSA and palmitic acid^44,45^), we also repeated APD studies on WTC MPS treated with MM in which we independently varied levels of these two components. The complete absence of palmitic acid and reduction of BSA levels to those provided by B-27 alone from MM led to APD that was not significantly different from APD observed for SM treated MPS (M1; **Fig. S3G**). In the absence of palmitic acid, the addition of a high dose of BSA appeared to be somewhat toxic to cardiac tissues, as the prevalence of beating in these samples (M2) was nearly zero, making it difficult to interpret the apparent APD (**Fig. S3H**). Interestingly, palmitic-acid enriched media still had significant effects on APD reduction, while negative effects of high albumin dosing were slightly reduced, when the total amount of albumin present was reduced below 2.5%. However, since the APD observed with 1% BSA (M4; **Fig. S3G**) was significantly lower than the APD for adult human left ventricular cardiomyocytes, we assumed these tissues might fall outside the ideal physiologic range for drug testing. Thus, MM with 125µM oleic acid, 100µM palmitic acid, 0.5g/l glucose, 10mM galactose and 2.5% BSA was used for all subsequent studies.

Although several types of fatty acids have been shown to enhance maturation of hiPSC-CM monolayers and engineered heart tissues, here we found that it was necessary to include palmitic acid specifically (**Fig. S3G**). This suggests that although generalized metabolic effects such as oxidation-induced DNA damage response^34^ are likely important to electrophysiological maturation of hiPSC-CM *in vitro*, events affected directly by palmitic acid, such as palmitoylation of calcium channel subunits^46^ or receptors^47^ may also be involved. Furthermore, although it was possible to reduce APD with maturation media that included oleic acid and palmitic acid without increased albumin (M4?; **Fig. S3G**), this treatment yielded APD that fell below the target range of adult cardiomyocytes. It is possible that albumin, which is a fatty-acid carrier *in vivo*, provided a more temporally stable dose of fatty acids to cells cultured within the MPS. The concentration of albumin in MM is not markedly dissimilar from the 3.5-5% albumin level found in human blood^48^. Finally, the finding that fatty-acid based maturation media had significant effects on action potential and pharmacology within WTC MPS, but not 2D hiPSC-CM monolayers, suggests the need to incorporate advanced 3D co-culture models during development of protocols to mature hiPSC-CM, and likely other hiPSC-derived tissue cells.

### Divergent Effects of Maturation Media on APD in Genetically Distinct hiPSC-Cardiomyocytes

We next tested fatty-acid based MM in a genetically distinct set of hiPSC-CM, SCVI20. In contrast to WTC, SCVI20 exhibited much shorter APD at baseline, ranging from 250 to 350ms in monolayers and 150 to 400ms in MPS (**Fig. 3C**). Treatment of SCVI20 MPS for 10 days with MM prolonged APD, so that APD_80_ fell within a very similar range to what we observed for MM-treated WTC MPS **(Fig. 3D,F)**, and within the range of 300 to 400msec observed for human adult ventricular cardiomyocytes (Brandenburger *et al*. 2012; ref. 49). Interestingly, analysis of Ca^2+^ handling using OGB-1-AM revealed that MM treatment produced a similar trend toward increased background normalized calcium amplitude (ΔF/F_0_) in SCVI20 MPS as it did in WTC (**Fig. 3I**). However, this trend was not statistically significant in SCVI20. Importantly, because GCaMP6f and OGB-1-AM have different affinity for Ca^2+^ and different mechanisms for Ca^2+^ induced fluorescence (*K*_*D*_ of 375nM for GCaMP^50^ and *K*_*D*_ of 170nM for OGB-1^51^), one cannot directly compare the ΔF/F_0_ response between MPS derived from these different lines. Despite not enhancing ΔF/F_0_ in SCVI20, MM did significantly increase the maximum Ca^2+^ upstroke velocity or MPS of this genotype (**Fig. 3J**). There was no trend between media type and upstroke timing in SCVI20 MPS (**Fig. S3B**).

The differences in cAPD_80_ of SCVI20 and WTC MPS in response to MM prompted us to test the effects of this media on a third cell line, SCVI273. This additional line responded similarly to WTC in terms of APD shortening effects of MM (**Fig. S3C**). However, there was no trend toward increased ΔF/F_0_ (**Fig S3D**). As with SCVI20, this lack of change in ΔF/F_0_ is likely to reflect the difference in calcium affinity of OGB-1 versus GCaMP6f^50,51^.

### Pre-Treatment of hiPSC-Derived-Cardiomyocyte Based MPS with Maturation Media Supports a Shift Toward a more Mature Metabolic Phenotype

Maturation-media induced changes in the state of mitochondria within iPSC-cardiomyocytes in MPS were next assessed. Mitochondrial inner transmembrane potential, as measured with MitoTracker Red, was markedly upregulated by MM in both WTC and SCIV20 (**Fig. 4**). This suggests an increase in oxidative phosphorylation^52^ in MM treated MPS. Antibody-staining revealed that overall mitochondrial density did not change, but mitochondrial organization shifted with MM treatment in both genotypes (**Fig. 4A-D**). SM produced short filaments and rounded structures, whereas MM treatment yielded a mitochondrial structure of extended filaments and networks. Similar filamentous networks of mitochondria, running perpendicular to Z-lines, have been observed previously during postnatal development of rodent cardiomyocytes^53^. Interestingly, although iPSC-cardiomyocyte monolayers treated with MM showed upregulation of maximum Oxygen Consumption Rate (OCR) in Seahorse assays, they did not show marked increased in mitochondrial membrane potential (**Fig. 4E,F, Fig. S4**). These data indicate that fatty acid-based MM directly alters the metabolic state of iPSC-cardiomyocytes, as has been shown in previous studies^34,35^. The finding that MM caused more substantial shifts in mitochondrial state in MPS as opposed to monolayer culture is consistent with our finding that MM caused much less significant changes in physiology of monolayers as compared to MPS (**Fig. 3**). This may reflect a lower ATP consumption rate in monolayers, which are not doing active mechanical work, as contrasted to cardiomyocytes in MPS that contract against PDMS posts.

**Figure 4.**
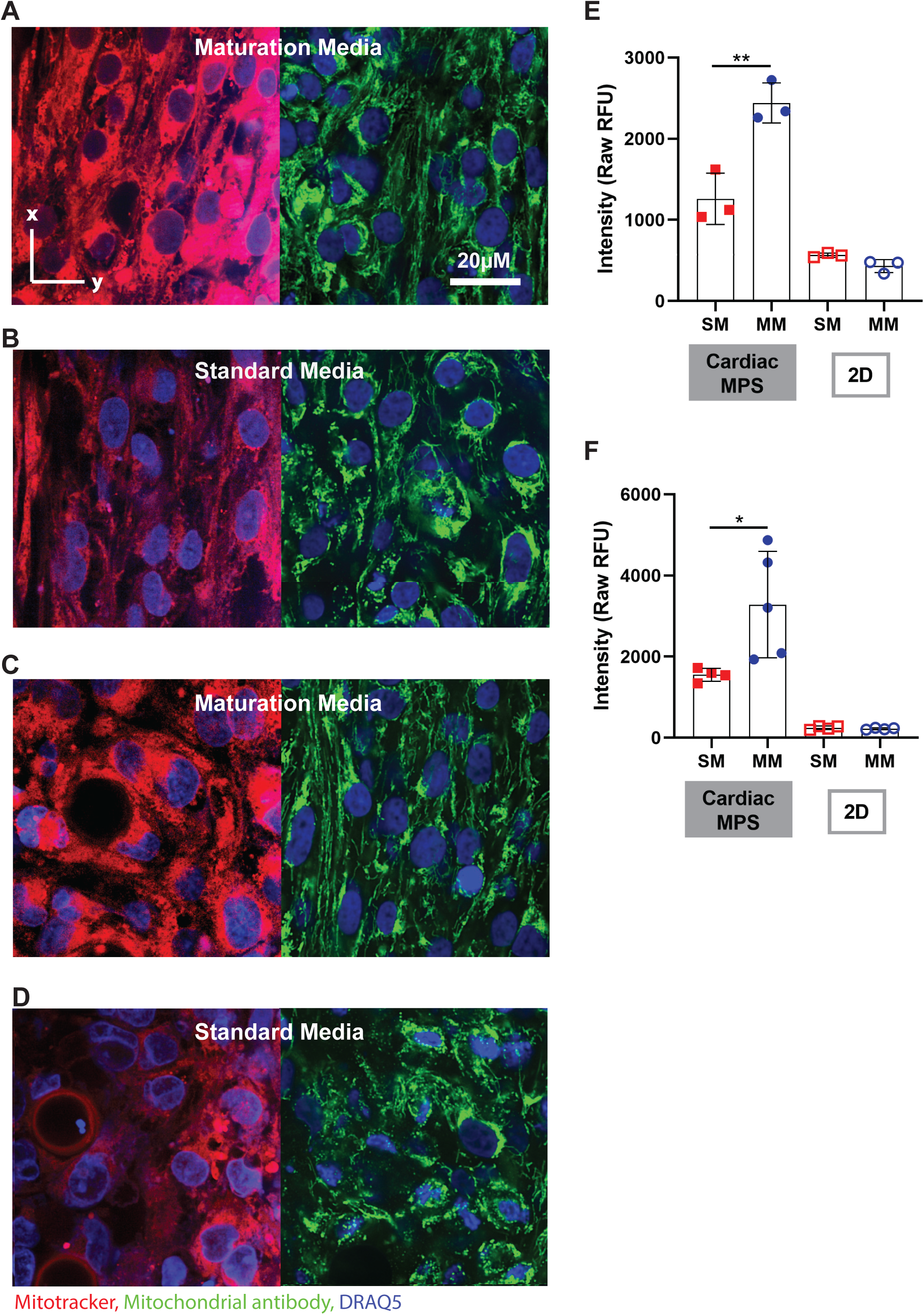
Metabolic Phenotype of MPS Treated with Maturation Media. **A-D)** Representative micrographs of (**A,B**) WTC and (**C,D**) SCVI20 MPS treated for 10 days with either (**A,C**) MM or (**B,D**) SM, then stained with MitoTracker Red (red; left) or anti-mitochondrial antibodies (green; right). Nuclei are counterstained with Draq5 (blue). **E,F)** Quantification of MitoTracker Red intensity in cardiac MPS and 2D monolayers for WTC (**E**) or SCVI20 (**F**) (each data point represents one independent MPS or monolayer well; mean MPS intensity values were calculated by averaging five randomly selected fields). (*\* p* < 0.05, *\*\* p* < 0.01, 2-way *t*-test).

### Pre-Treatment of hiPSC-Derived-Cardiomyocyte Based MPS with Maturation Media Supports Inotropic Responsiveness

MM treatment did not result in changes in gross sarcomere structure within WTC or SCVI20 MPS, as assessed by staining for sarcomeric α-Actinin (ACTN2; **Fig. 5A-B**). Quantitative analysis of sarcomere morphology with Fourier-Transform based methods^43,54^ was consistent with these qualitative observations, and suggested no substantial changes in sarcomere organization as a result of MM treatment (**Fig. S5A-B**). Expression and localization of β-Myosin Heavy Chain and MLC-2v, two sarcomere-associated proteins which typically mark mature, ventricular-like iPSC-cardiomyocytes, were very similar in MM and SM pre-treated MPS (**Fig. S5C-G**). There was a trend toward upregulation of MLC-2v protein in SCVI20 MPS, consistent with previous observations by Mills *et al*. that palmitic-acid enriched, glucose depleted media increased MLC-2v levels^34^.

**Figure 5.**
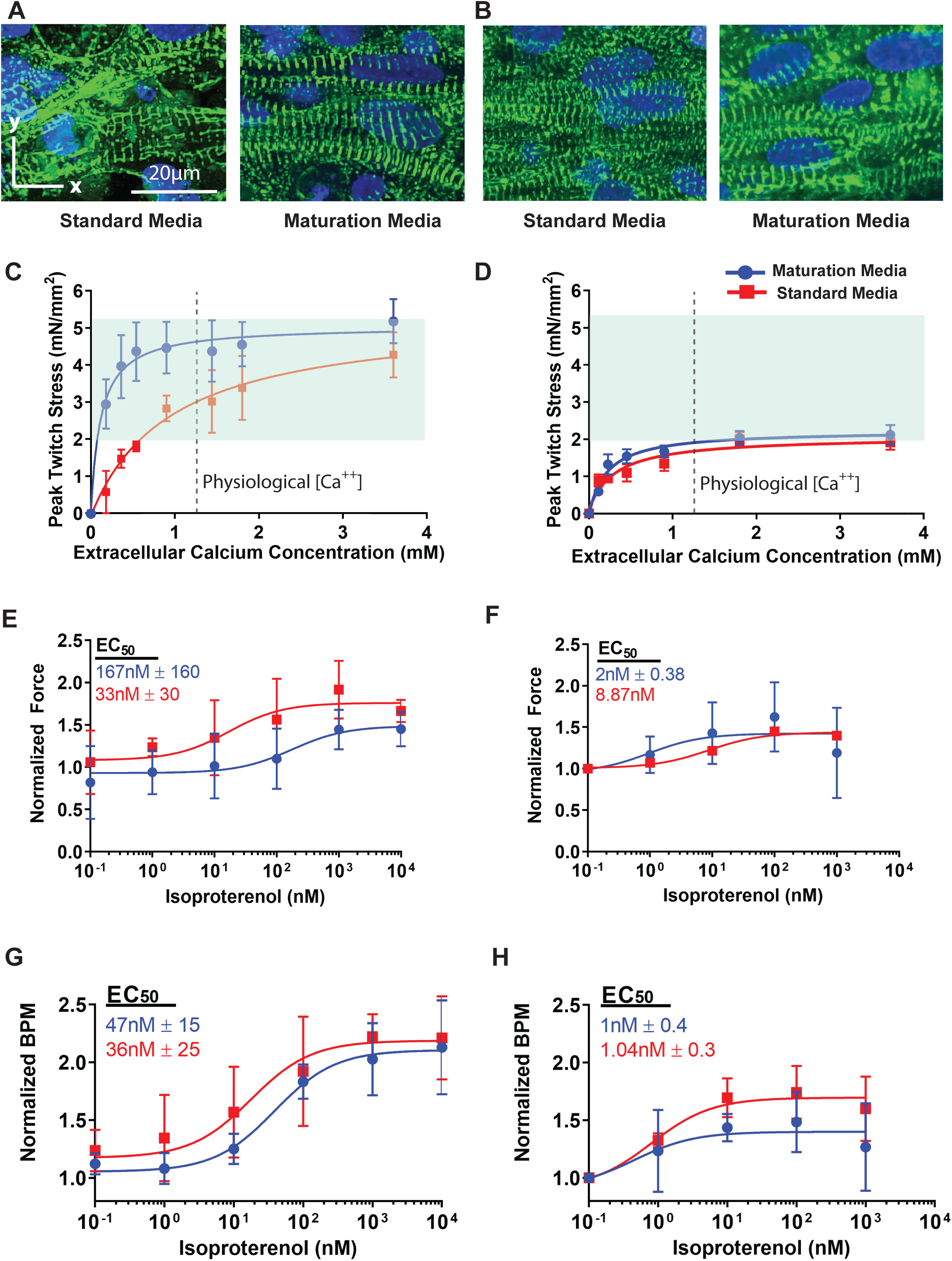
Inotropic Responsive of Maturation Media Treated MPS. **A,B)**Representative confocal micrographs depicting sarcomere morphology (Sarcomeric α-Actinin Staining, green, with DAPI nuclear counterstain, blue) of MPS treated for ten days with either standard media (SM) or maturation media (MM) for **A)** WTC and **B)** SCVI20. Contractile stress generated by MPS pre-treated with either SM or MM as a function of extracellular calcium (delivered in Tyrode’s saline) for **C)** WTC and **D)** SCVI20. The translucent green box denotes the force generated by adult human heart slice cultures (Brandenburger *et al*. 2012). Normalized (**E**) force and (**G**) beat-rate as a function of isoproterenol dose in SM and MM pre-treated WTC MPS. Normalized (**F**) force and (**H**) beat-rate as a function of isoproterenol dose in SM and MM pre-treated SCVI20 MPS. For force calculation (inotropy), MPS were cultured in Tyrodde’s saline with 0.9mM Ca^2+^. For beat rate calculation (chronotropy), MPS were first equilibrated to standard media and then received isoproterenol doses in this media. Data: mean ± *SEM, n* = 3-5. Scale bars: **A**: left panels, 20µm and right panels, 10µm.

When we measured force developed in MPS via the deflection of PDMS pillars in the chamber (**Supplemental Methods; Fig. S6)**, we found that both SM and MM-pre-treated WTC MPS exhibited a robust dose response to increased extracellular calcium, with a maximal range of stress similar to adult human heart tissue slices (**Fig. 5C**; data on adult slices calculated from Brandenburger *et al*.; ref. 49). Consistent with work by Mills, we observed that fatty-acid based maturation media neither inhibited nor enhanced peak twitch force^34^. However, MM pre-treated WTC MPS were sensitized to lower concentrations of extracellular calcium than SM pre-treated controls, with a statistically significant EC_50_ of 0.11±0.06 mM for MM pre-treated MPS versus 0.95±0.46mM for SM-pre-treated controls (*p* < 0.05, 2-way *t*-test). Furthermore, MM pre-treated MPS showed a statistically significant steeper fold-increase in force in response to extracellular calcium over the linear region of the calcium-force response curve, with an initial slope of 12±1.5 mN/mm^2^/mM Ca^2+^ versus 3.9±0.26 for SM-pre-treated controls (*p* < 0.01, 2-way *t*-test). Consistent with the trend that MM-treatment enhanced Ca^2+^ handling (ΔF/F_0_) more significantly in WTC than in SCVI20 MPS, SCVI20 MPS also showed a trend toward increased force with MM, but even at low extracellular Ca^2+^ levels, the trend towards higher twitch forces with MM was not statistically significant **(Fig. 5D)**.

Analysis of tissue inotropic responses to isoproterenol in MM pre-treated WTC MPS showed a trend toward desensitization, and MM-treated SCVI20 MPS showed a trend toward sensitization to this drug, though neither trend was statistically significant (**Fig. 5E,F**). Similarly to inotropic effects of isoproterenol assessed at constant (paced at 1Hz) beat-rate, MM pre-treatment appeared to slightly desensitize MPS to the chronotropic effects of isoproterenol in both genotypes, although, as with inotropic effects, the changes were not statistically significant (**Fig. 5G,H**). The EC_50_ values for isoproterenol chronotropy fell within the range recently reported for engineered heart tissue subjected to exercise-induced maturation via external pacing^23^, and within the range of adult human heart slices^49^.

Collectively, these data suggested that MM does not damage sarcomeres or interfere with excitation-contraction coupling or adrenergic responsiveness and enhances calcium contraction coupling when the amount of extracellular calcium is limiting. Furthermore, unlike the divergent trend observed with respect to MM-induced change in APD in SCVI20 vs. WTC, inotropic responsiveness showed a relatively similar trend in both genotypes, suggesting multimodal analysis of tissue function for accurate characterization.

### Gene Expression Changes caused by Maturation Media in MPS-Cultured hiPSC-CM

Consistent with previous studies, we observed genotype-based variability in transcription of monolayer hiPSC-CM in standard media^11^. However, there was no significant difference in a panel of ion channel and sarcomere related genes caused by MM treatment of monolayers of either genotype **(Fig. 6A,B)**. Consistent with observations regarding the immaturity of hiPSC-CM, several ion channel transcripts were either deficient or overexpressed in these cells, when compared to commercially available RNA obtained from adult human hearts (**Fig. 6A,B**). Further, the absolute level of SCN5A (related to sodium current, *I*_*Na*_) and several other ion channel transcripts, including KCNJ2 (related to *I*_*K1*_), was highly variable between different batches of purified, 2D monolayer hiPSC-CM. This may point to mis-regulation of expression in these channels in non-physiologic culture formats, or to differences in the relative levels of different cardiomyocyte sub-types obtained from our small molecule-based differentiation protocol^55,56^.

**Figure 6.**
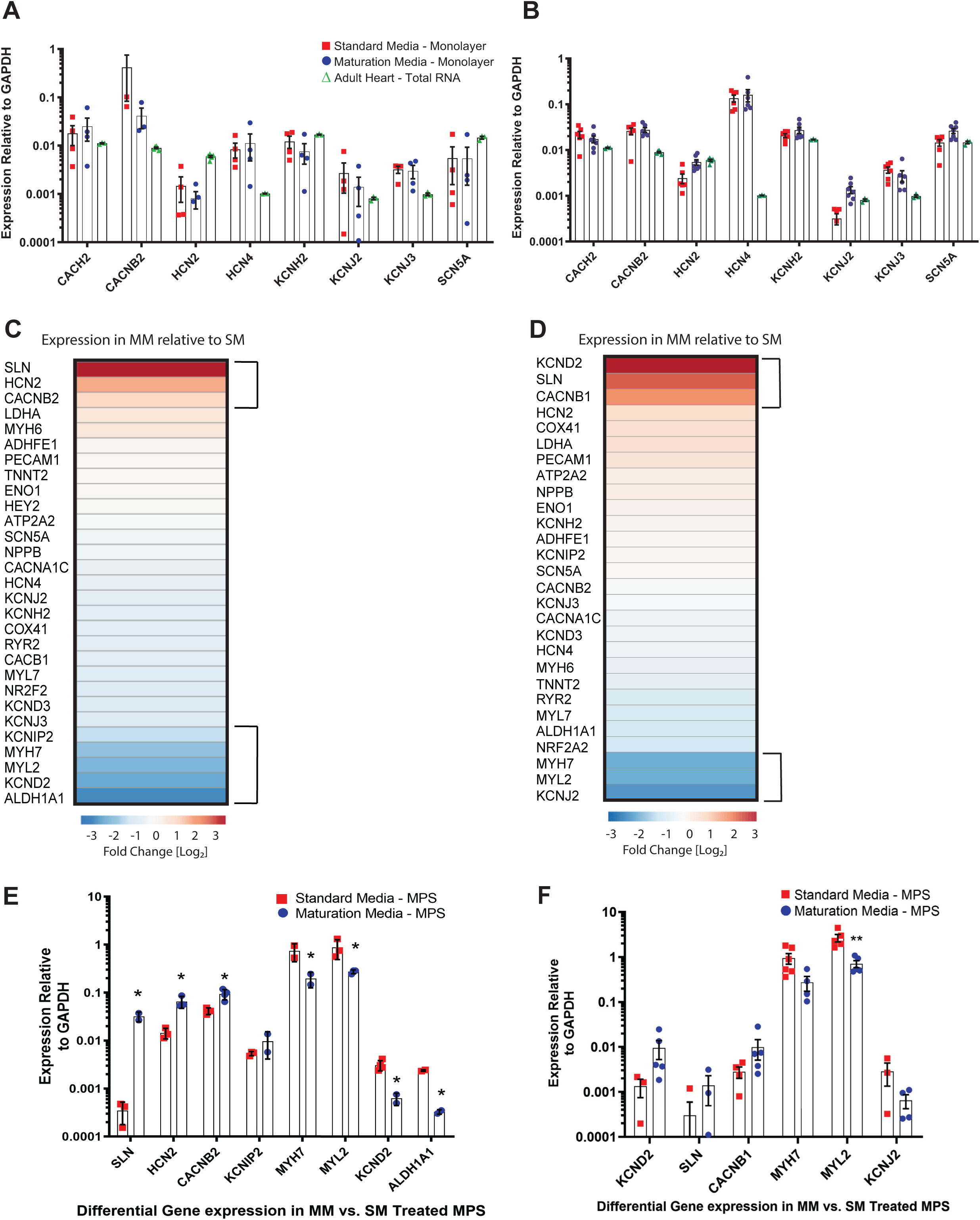
Gene Expression analysis of Monolayers and MPS Treated with Lipid-Based Maturation Media. Quantitative RT-PCR analysis of expression of ion channel and sarcomere transcripts in hiPSC-CM 2D monolayers after ten days of culture in either Standard (SM; red) or Maturation Media (MM; blue) for **A)** WTC and **B)** SCVI20. None of the genes tested exhibited statistically significant expression changes as a result of Maturation Media. Data: plot of points with median, *n* = 4. Heat-map of relative gene expression in MM-treated MPS as compared to SM-treated MPS, as assessed by qRT-PCR on cDNA libraries amplified from RNA isolated of MPS treated for ten days with SM or MM for **C)** WTC and **D)** SCVI20. Specific analysis for **E)** WTC and **F)** SCVI20, indicating individual biological replicates (treatments of hiPSC-CM obtained from independent differentiations) of differentially expressed transcripts for ion channels or sarcomere genes in MM and SM treated MPS. MPS PCR data were plotted on ClustVis to obtain heatmaps of the gene expression. The genes within 70% percentile of differential expression were then selected and plotted (**C,D**). Error bars: SEM, *n* = 4. * *p* < 0.05, ** *p* < 0.01, 2-way t-test, with Mann-Whitney correction method.

We next assessed gene expression in MPS treated for ten days with either SM or MM. Interestingly, and in contrast to results by Mills *et al*.^*34*^, we did not observe significant variation in expression of the glycolysis associated gene, GAPDH, relative to other potential “housekeeping” genes (data not shown). A lack of change in GAPDH expression on the protein level was further verified by immunostaining, which revealed no significant effects of MM on GAPDH expression in either WTC or SCVI20 (**Fig. S7**).

Analysis of a panel of genes involved in electrophysiology, cell identity, contractility and calcium handling did not reveal a global shift in expression as would be expected for gross changes in cell differentiation or population composition in either genotype (**Fig. 6C,D**). Further, we did not observe substantial differences in the transcript expression for most potassium channels, including hERG/KCNH2 (related to *I*_*Kr*_). However, MM-treated WTC MPS showed upregulation of SLN, HCN2 and CACNB2, and downregulation of KCND2 and ALDH1A1 (**Fig. 6E**). In SCVI20, MM treatment caused a trend toward decreased KCNJ2; this gene is related to *I*_*K1*_, and diminishing this current could be one explanation for increasing APD. We also observed KCND2 (*I*_*to*_) upregulation, which increases robustness by counteracting KCNJ2 decrease. When treated with MM, SCVI20 MPS also showed upregulation of CACNB1 and SLN, although these changes were not statistically significant **(Fig. 6F)**.

Sarcolipin (SLN) is known to bind the SERCA pump in the sarcoplasmic reticulum of cardiac and skeletal muscle and act as a regulatory protein to increase heat production^57^. It does so by partially uncoupling SERCA’s calcium re-sequestration function from ATP hydrolysis and therefore burn extra ATP without influencing calcium transients^58–60^. It is possible that since MM brings a higher caloric intake to the tissue through fatty acids, MM-treated MPS tissues might upregulate SLN to burn excess ATP. This would be consistent with more mature and efficient metabolism in MM through a metabolic switch from glycolysis to oxidative phosphorylation, which is supported by direct analysis of mitochondrial morphology and transmembrane gradients in MM-treated MPS (**Fig. 4**). SLN is preferentially expressed in atria and in previous work has been used to identify atrial-like iPSC-cardiomyocytes^61^. However, the possibility that MM shifted cardiomyocytes in MPS toward atrial subtypes is contracted by the overall pattern of gene expression – specifically, transcript levels for HEY2, a ventricular cardiomyocyte-specific transcription factor^62^, were not affected by MM in WTC (**Fig. 6C**). Similarly, atrial cardiomyocyte specific transcription factor NR2F2^62^ and sarcomere gene MYL7 were both downregulated by MM. Altogether, these data suggest that upregulation of SLN is more likely to be directly related to cardiomyocyte metabolism than tied to cardiomyocyte subtype.

The β-subunits of the voltage activated calcium channel expression encoded by CACNB1 and CACNB2 are required for expression of the *L*-type calcium current through regulating trafficking and activation of α-subunits^46,63^. Changes in the levels of these subunits could potentially alter *I*_*CaL*_. Despite our observation that force production and inotropic responsiveness of MPS were not perturbed by MM pre-treatment, we observed significant downregulation of several genes associated with calcium handling and sarcomere function in WTC, including MYL7, MYL2/MLC2V, and MYH7 (**Fig. 6E,F**). We observed a similar reduction of MYL2 and a trend towards decreased MYH7 in SCVI20. However, our data from immunostaining suggests that levels of the proteins encoded by the genes are not changed significantly, and trend toward upregulation with MM, in both genotypes (**Fig. S5**), with no changes in cellular localization. This finding suggests that gene expression data does not necessarily correlate perfectly with protein level and localization. However, despite relatively similar levels of MYL2 and MYH7 proteins in WTC and SCVI20, the contractile forces of WTC are substantially higher (**Fig. 5C,D**).

### Mathematical modeling of the ion channel and calcium transient contribution to electrophysiological patterns in MM-treated MPS

Although MM has divergent effects on APD in SCVI20 vs. WTC MPS, in both cases, this fatty acid-based media brings tissues closer to an *in-vivo*-like ventricular APD. To gain insight into how this occurs, we applied newly developed mathematical models^64^ to estimate the contribution of individual Na^+^, K^+^, and Ca^2+^ currents, along with calcium handling machinery, to the experimentally measured action potential and calcium transients (**Fig. 7A-C**).

**Figure 7.**
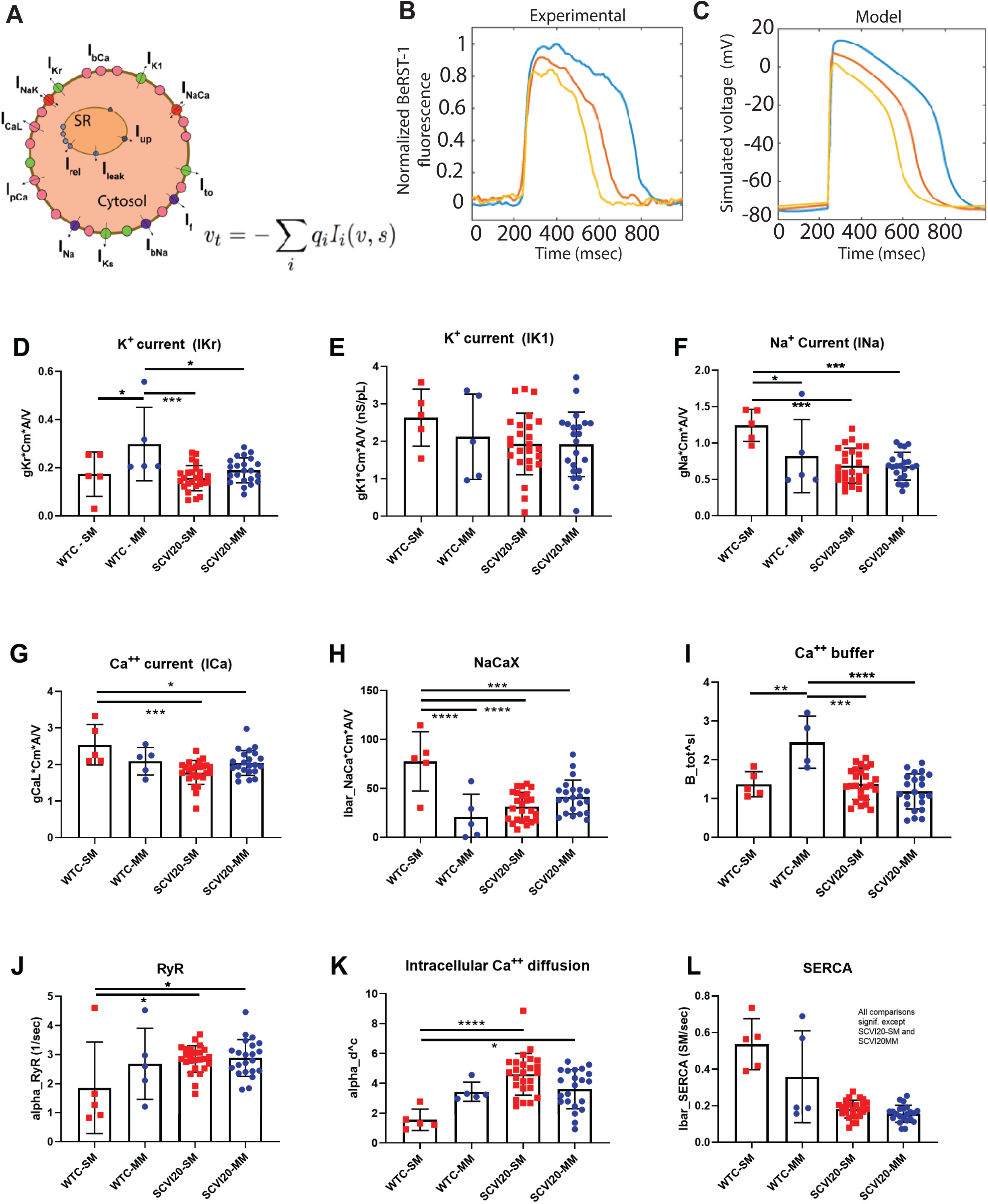
Mathematical Modeling of the Contribution of Individual Currents and Calcium Handling Machinery to the Action Potential of Monolayers and MPS. **A)** Schematic of the model. **B)** Examples of experimentally measured individual currents (data obtained with BeRST-1) used as model inputs. **C)** Representative simulated currents based on the corresponding color of experimentally measured current. **D-G)** Simulated current fluxes. **D-E)** Major potassium currents: D, *I*_*K1*_ and E, *I*_*Kr*_ (hERG). **F)** Sodium current, *I*_*Na*_. **G)** *L*-type Calcium current, *I*_*CaL*_. **H)** Sodium-calcium exchange current, *I*_*NaCax*_. **I-L)** Simulated calcium dynamics. **I)** Calcium buffer; **J)** Ryanodine-receptor flux. **K)** Intracellular Ca^2+^ diffusion. **L)** SERCA pump activity. Media type and genotype had global effects (1-way ANOVA) in all panels except E. * *p* < 0.05, *\*\* p* < 0.01, *p* < 10^−3^, post-hoc Tukey tests.

In contrast to gene expression data, but consistent with observed changed in APD, simulations predicted enhanced conductance (g_x_•C_m_•A/V) for the late rectifier potassium current *I*_*Kr*_ (hERG) in WTC (**Fig. 7D**). There was also a trend, albeit not statistically significant, toward increased *I*_*Kr*_ in SCVI20 MPS with MM treatment (**Fig. 7D**). No trends were observed in predicted *I*_*K1*_ **(Fig. 7E)**. In WTC MPS, simulations also predicted significant reduction of Na^+^ current, enhancement of intracellular Ca^2+^ buffering, and repression of sodium-calcium exchange current (*I*_*NaCaX*_) with MM (**Fig. 7F,H-I**); they also predicted trends toward reduced *L*-type Ca^2+^ current and improvements in intracellular calcium transport and RYR function with MM (**Fig. 7G,J,K**). In SCVI20 MPS, MM-treatment did not result in significant changes in any of the simulated currents or calcium handling parameters described by the model. However, the trend toward increased *I*_*Na*_, *I*_*CaL*_, *I*_*NaCx*_ and diminished intracellular calcium transport with MM treatment would be expected to delay the onset of repolarizing currents.

Simulations predicted diminished SERCA pump activity in MM-treated MPS of both genotypes; this trend was significant in WTC **(Fig. 7L)**. These results are consistent with gene expression data predicting marked MM-induced upregulation of SLN in WTC MPS, and a trend toward upregulation of this gene in SCVI20 MPS. SLN interacts with and suppresses the SERCA pump.

Consistent with the ΔF/F_0_ response (**Fig. 3G**) and the force developed in response to low extracellular calcium levels (**Fig. 5C**), the model shows a significant shift towards higher intracellular calcium in MM-WTC compared to SM-WTC. The overall calcium balance and SR stocks at steady state is shifted towards influx. Lower NCX activity and increase in Ca diffusion in MM-WTC (**Fig. 7H,K**) will rapidly bring calcium away from *I*_*CaL*_ and NCX towards sarcomere, acting as a buffer (**Fig. 7I**). Overall, the calcium accumulation in the cell will limit the inward currents and participate in APD shortening. In the case of MM-SCVI20, although the trend showed an increase of ΔF/F_0_ (**Fig. 3I**), it was not significant compared to SM-SCVI20. This is consistent with the model showing a trend towards an increase in *I*_*CaL*_ and an increase in NCX (**Fig. 7G,H**) which could help explain the increase in APD and the constant calcium transient.

Consistent with the markedly different action potential waveforms in SCVI20 versus WTC MPS in standard media, simulations predicted significant differences in almost every ion current and calcium handling parameter described by the model (except for potassium currents and calcium buffering; **Fig. 7D-E,J**) for SCVI vs. WTC MPS. Interestingly, and consistent with our observation that fatty acid media brought APD to a very similar range for both genotypes, MM-treatment led to nearly identical predicted currents and calcium handling: no parameters predicted by the model differ between MM-treated SCVI20 and WTC MPS, except for *I*_*Kr*_ and calcium buffering, which were predicted to be higher in MM-treated WTC MPS **(Fig. 7D,I)**. The observation that no single simulated current was significantly different in MM vs. SM treated SCVI20 MPS suggests the potential that SCVI20 hiPSC-CM emerge from cardiac differentiation in a more matured state than do WTC hiPSC-CM. However, the potential that SCVI20 hiPSC-CM are more mature is contradicted by the significantly higher force produced by WTC MPS in either SM or MM (**Fig. 5C,D**). It is more likely the case that the significant changes in APD_80_ between SM and MM treated SCVI20 MPS (**Fig. 3D,E,H**) are caused by concurrent but subtle changes in several of the channel fluxes that contribute to the action potential.

To test the possibility that MM treatment causes more calcium to be stored in the sarcoplasmic reticulum of hiPSC-CM, we pharmacologically perturbed SR calcium handling in SM and MM pre-treated WTC MPS. Normally, in mature cardiomyocytes, Ca^2+^ released from the SR via RyR potently inactivates *I*_*CaL*_ and curtails sarcolemmal Ca^2+^ influx^65^. However, inhibition of the SERCA pump reduces the SR store, which in turn reduces systolic SR Ca^2+^ release and calcium-dependent inactivation of the membrane-bound L-type calcium channels. This mechanism is supported by observations of MPS treated with a saturating (10mM) dose of Ryanodine, which blocks the cardiac ryanodine receptor. This treatment dramatically increased both Ca^2+^ rise-time and decay time (τ_75_) in MM-pre-treated, but not SM pre-treated WTC MPS. Moreover, although Ryanodine diminished ΔF/F_0_ for both SM and MM pre-treated MPS, MM pre-treated MPS showed a more substantial change in ΔF/F_0_ in response to this drug (**Fig. S8**). Neither MM nor SM pre-treated MPS displayed a change in beat rate upon treatment with Ryanodine (**Fig. S8A**). Not only are these findings consistent with predictions from simulations of trend toward greater predicted steady state SR-storage of Ca^2+^ with MM (**Fig. 7**), they are also consistent with a trend toward greater RyR flux in MM treated WTC MPS (**Fig. 7J**).

A second drug, Thapsigargin, which blocks SR Ca^2+^ reuptake via SERCA, also markedly slowed Ca^2+^ reuptake, increasing τ_75_, in MM but not SM-pretreated MPS. In contrast to Ryanodine, Thapsigargin did not affect ΔF/F_0_ or calcium rise-time (**Fig. S9**). This response is consistent with our simulation data, which suggest that although SR calcium storage is enhanced by fatty acid media, there is some predicted suppression of SERCA activity (**Fig. 7L**). Altogether, these experimental results are consistent with the prediction of the simulations that pre-treatment with MM increases the SR Ca^2+^ store, and its contribution to the cytosolic Ca^2+^ transient that fuels contraction. These results are internally consistent with the results of our simulations, and with previous work where matured hiPSC-CM exhibited a more marked calcium decay time in response to thapsigargin^36^. In previous studies, maturation media containing dexamethasone and triiodothyronine^66^, and maturation induced by continuous field pacing^23^ have been linked to improvements in hiPSC-CM Ca^2+^ handling through formation of T-tubules. However, in the present study, we observed minimal evidence of T-tubules, although qualitative analysis of Wheat-Germ Agglutinin stained MPS showed some instances of membrane invaginations that were more prominent in MM (**Fig. S10A-C**) than in SM (**Fig. S10D,E**) treated MPS.

### Pre-Treatment of hiPSC-Derived-Cardiomyocyte Based MPS with Maturation Media Enhances Prediction of Drug Induced Pro-Arrhythmia

We next assessed whether maturation media treatment could lead to more predictive pharmacology of compounds with known pro-arrhythmic effects, and further whether the simulations predicted pharmacology changes induced by MM-pre-treatment. Pro-arrhythmia effects of these drugs have been quantified in clinical studies by measuring the QT interval of the Electrocardiogram (ECG)^67^. One important drug from these studies is Verapamil, which blocks both *I*_*CaL*_ and *I*_*Kr*_, is routinely used in the clinic to shorten the QT interval in patients prone to suffer arrhythmias based on prolonged APD (for example, patients with long-QT syndrome^67^) based on *I*_*CaL*_ block. Verapamil is a prototypical example of a “false positive” drug that would appear to prolong APD and QT interval if one only assessed block of the current encoded by hERG/KCNH2, *I*_*Kr*_^67,68^.

When we analyzed Verapamil dose-escalation effects in field-paced MPS and used beating prevalence as the metric for characterizing IC_50_, we observed that MM-pre-treated WTC MPS exhibited enhanced Verapamil resistance compared to SM-treated MPS (971 nM for MM-pre-treated MPS, versus 90nM for SM-treated MPS; **Fig. 8A**). In contrast, SCVI20 MPS showed a decreased Verapamil resistance with the same media (**Fig. 9A**). Direct analysis of APD revealed a dose-dependent decrease in APD_80_, consistent with the clinical application of this drug to shorten QT duration (**Fig. 8B, 9B**). However, unlike beating prevalence, the dose-response for APD_80_ did not change appreciably in MM pre-treated, compared to SM-pre-treated MPS, for either genotype. There were, however, weak trends toward Verapamil desensitization in MM-treated WTC and Verapamil sensitization in MM-treated SCVI20 MPS when APD_80_ was used as an *in vitro* metric of drug response (**Fig 8C, 9C**).

**Figure 8.**
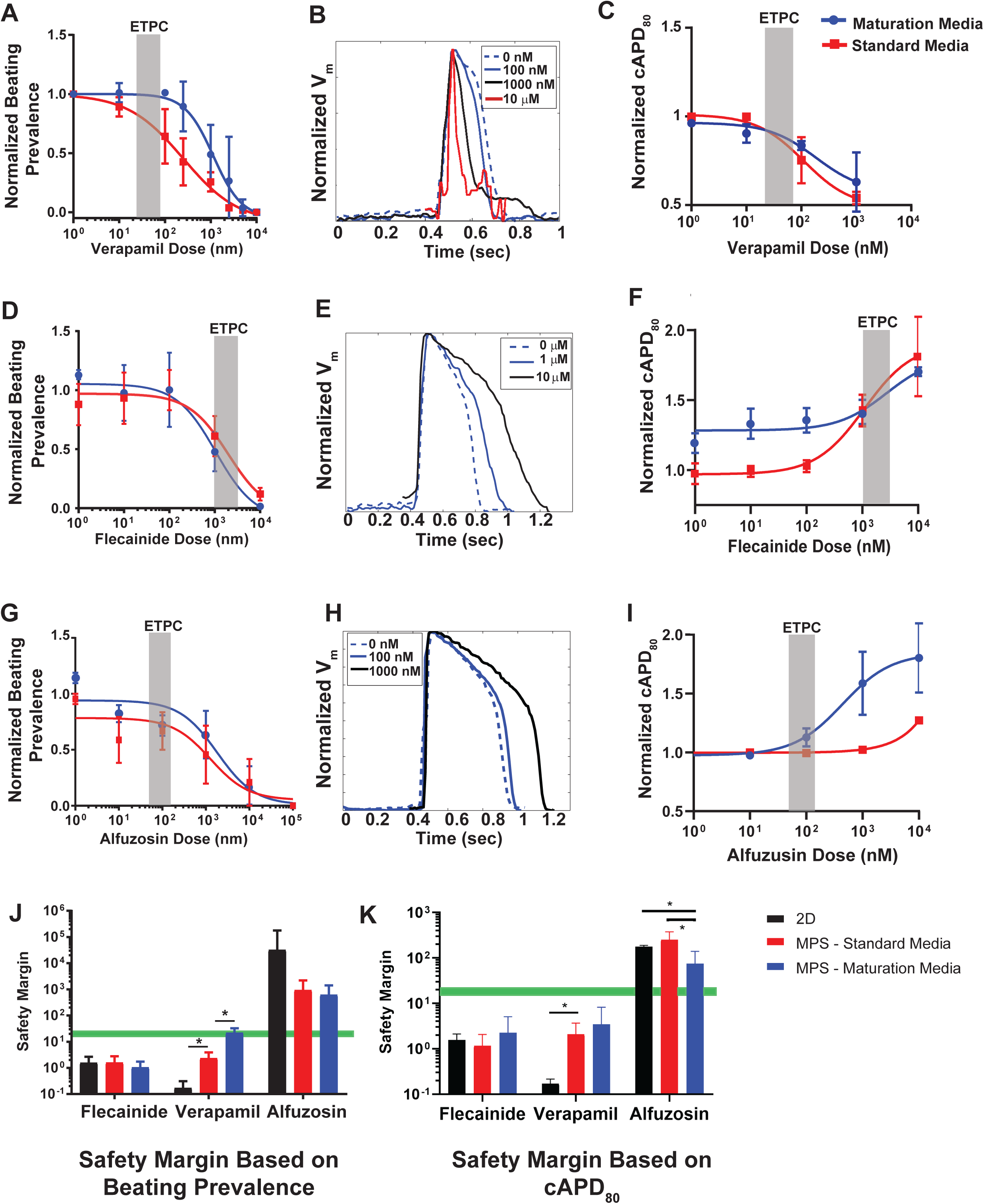
Proarrhythmia Pharmacology of Matured WTC Cardiac MPS. IC_50_ and EC_50_ analyses were performed in MPS pretreated for ten days with either Maturation Media (blue curves) or Standard Media (Red Curves). For all studies, MPS were equilibrated to Standard Media, and then exposed to escalating doses of **A-C)** Verapamil, **D-F)** Flecainide, and **G-I)** Alfuzosin. IC_50_ curves were obtained by measuring beating prevalence (**A,D,G**) and EC_50_ curves by measuring 80% Action Potential Duration (APD_80_; **C,F,I**). Representative, intensity normalized action potential traces are depicted for MM-pretreated MPS for each drug (**B,E,H**). Estimated Therapeutic Plasma Concentration (ETPC) values were obtained from the literature. **J-K)** Safety margins (the ratio of *in vitro* IC_50_ from prevalence measurements or EC_50_ from APD_80_ measurements to literature values for ETPC) calculated based on **J**) beating prevalence and **K**) APD_80_. All MPS were paced at 1 Hz for pharmacology analysis. Data: mean ± *SEM, n* = 3-5. (* *p* < 0.05, 2-way *t-*test with Holm-Bonferroni correction for multiple comparisons).

**Figure 9.**
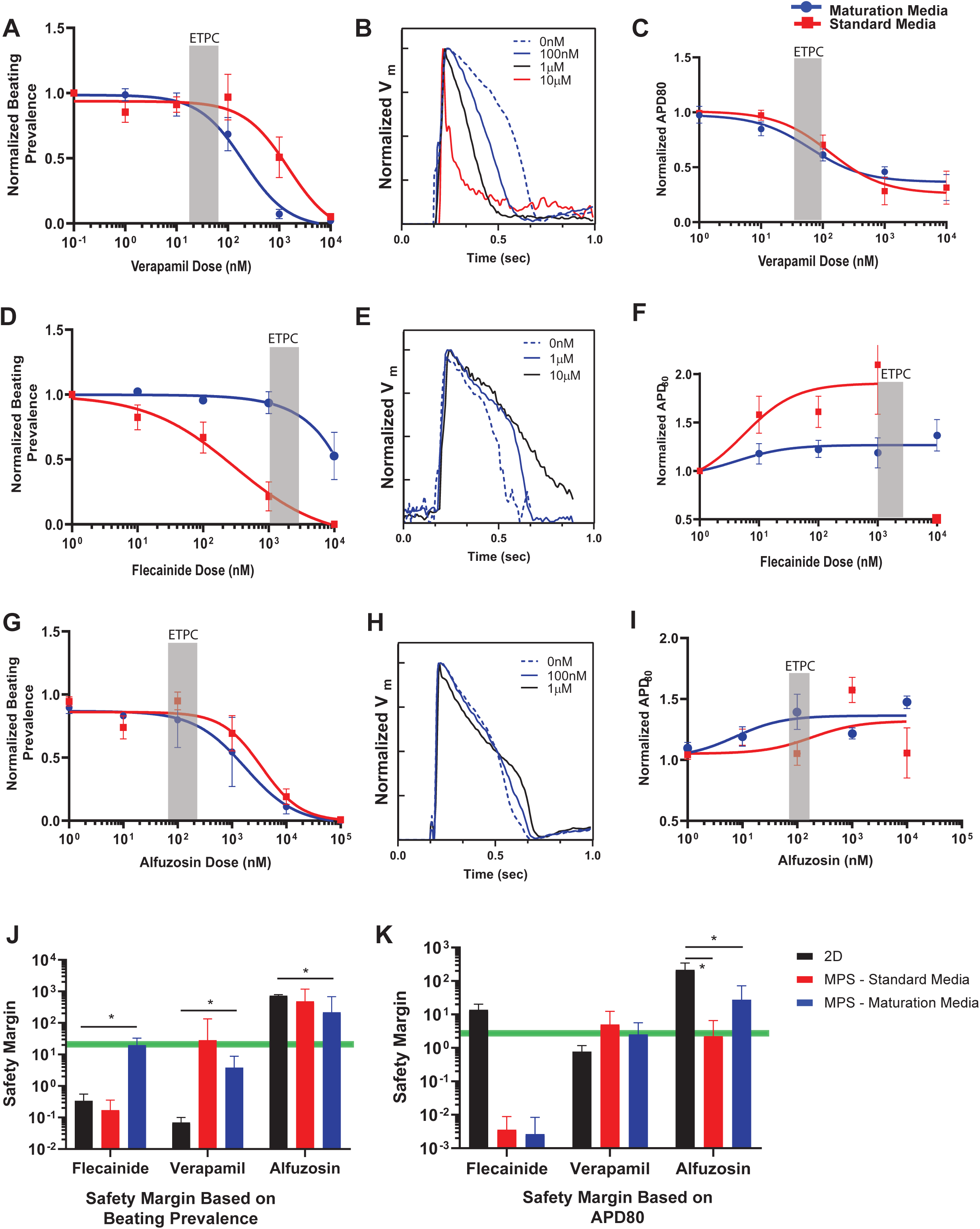
Proarrhythmia Pharmacology of Matured SCVI20 Cardiac MPS. IC_50_ and EC_50_ analyses were performed in MPS pretreated for ten days with either Maturation Media (blue curves) or Standard Media (Red Curves). For all studies, MPS were equilibrated to Standard Media, and then exposed to escalating doses of **A-C)** Verapamil, **D-F)** Flecainide, and **G-I)** Alfuzosin. IC_50_ curves were obtained by measuring beating prevalence (**A,D,G**) or 80% Action Potential Duration (APD_80_; **C,F,I**). Representative, intensity normalized action potential traces are depicted for MM-pretreated MPS for each drug (**B,E,H**). Estimated Therapeutic Plasma Concentration (ETPC) values were obtained from the literature. **J-K)** Safety margins (the ratio of *in vitro* IC_50_ or EC_50_ to literature values for ETPC) calculated based on **J**) beating prevalence and **K**) APD_80_. All MPS were paced at 1 Hz for pharmacology analysis. Data: mean ± *SEM, n* = 3-5. (* *p* < 0.05, 2-way *t-*test with Holm-Bonferroni correction for multiple comparisons).

These genotype-based shifts in Verapamil responsiveness may be explained by the relative *I*_*CaL*_ and *I*_*NaCax*_ levels predicted by mathematical models **(Fig. 7G,H)**: in WTC, the trend with MM is toward slightly higher *I*_*CaL*_ (increases intracellular Ca^2+^) and markedly lower *I*_*NaCax*_ (decreases intracellular Ca^2+^). The net effect of this increases influx and diminished efflux would be higher steady state levels of Ca^2+^ stored in the sarcoplasmic reticulum (SR). Higher SR-calcium levels would tend to increase contractility in response to Ca^2+^ uptake through *L*-type calcium channels, thereby desensitizing cells to *I*_*CaL*_ block. Increased steady state Ca^2+^ levels would also be consistent with the observed increase in calcium amplitude (ΔF/F_0_; **Fig. 3G**) and contractility in MM-WTC relative to SM-WTC MPS, in the setting of limiting extracellular calcium (**Fig. 5C**). In contrast, within SCVI20 MPS, the trend toward a stronger increase in *I*_*NaCax*_ than in *I*_*CaL*_ would result in lower SR levels of stored Ca^2+^. This would likely make cells more sensitive to *I*_*CaL*_ block.

We next assessed the ability to predict pharmacology of Flecainide, a class Ic (Na^+^ channel blocker) antiarrhythmic drug typically used to treat tachy-arrythmia, has been noted to have a narrow therapeutic index and is counter-indicated in patients with pre-existing structural disease^69^. This drug also blocks the hERG current^70^. Consistent with this narrow therapeutic index, we correctly observed very little difference between the Estimated Therapeutic Plasma Concentration (ETPC) of 1.5µM, and *in vitro* EC_50_ (APD_80_) and IC_50_ (beating prevalence) (**Fig. 8D-F, Fig. 9D,F**). Unlike Verapamil, Flecainide did not exhibit a differential EC_50_ within MM versus standard media pre-treated MPS for the WTC genotype; however, there was a weak trend toward desensitization of MM-pre-treated WTC MPS to the APD prolonging effects of this drug, which is consistent with improved hERG current in MM-treated WTC MPS. In the SCVI20 genotype, we observed a marked desensitization of MM-treated MPS to both contractile and APD changes induced by Flecainide. Although MM increases APD in SCVI20 MPS, the fatty acid media still causes a trend (albeit, not statistically significant) toward increased hERG current (**Fig. 7D**). Improved hERG current would desensitize MM-treated SCVI20 MPS toward *I*_*Kr*_ blocking effects of Flecainide.

Given how sensitive the SM-treated SCVI20 MPS are to the AP altering effects of Flecainide (EC_50_ of 5.5nM), the depressed contractility at 1 Hz pacing may be due to impaired calcium uptake. To test this hypothesis, we compared Ca^2+^ amplitude using OGB-1-AM imaging. This analysis revealed that, concurrent with lengthening the calcium transient timing, Ca^2+^ amplitude increased almost 3-fold over baseline in MM-SCVI20 MPS treated with 1μM Flecainide. In contrast, the amplitude of Ca^2+^ did not exceed baseline in Flecainide treated SM-SCVI20-MPS (**Fig. S11A-C**). The sodium-calcium exchanger can operate in reverse during action potential prolongation^65^. Thus, the observed ability of MM-SCVI20 MPS to withstand the contractility-depressing effects of Flecainide at doses that significantly disrupt contractility in SM-SCVI20-MPS may potentially be explained by the trend in simulations **(Fig. 7H)** toward enhanced NaCx exchange current in MM vs. SM pre-treated SCVI20 MPS. The observation that although Ca^2+^ uptake increases near 1μM Flecainide, but that contractility (beating prevalence) did not increase at this dose in MM-SCVI20 MPS, may potentially be explained by Flecainide-induced RYR block. Consistent with the exaggerated effects of flecainide on APD in SM-treated SCVI20 MPS, MPS of this genotype were more prone to Delayed After Depolarizations (DADs) that MM-treated MPS at 100nM dose of this drug (**Fig. S11D,E**).

Finally, we assessed MPS pharmacology of Alfuzosin, an α_1_-adrenergic blocking agent that has been shown to increase patients’ QT interval by hERG-independent mechanisms^11,71^. This makes Alfuzosin a “false negative” drug in screens for potential QT prolongation that rely on overexpression of the hERG/KCN2 gene in heterologous cell types. Here, we observed that in WTC, both MM and SM pretreated MPS exhibited IC_50_ near 1µM when measuring beating prevalence (**Fig. 8G**). However, when we tested the effects of this drug on extending APD_80_, we observed a specific sensitization with MM-pre-treated MPS, relative to MPS pre-treated with SM (**Fig. 8I**). We can explain the drastic change in APD with an unchanged prevalence through the efficient calcium transport of the MM-WTC, enabling the cells to contract despite the short resting time between long-duration actional potentials. In SCVI20, there was a trend toward sensitization of SM-treated MPS to Alfuzosin, although differences in the EC_50_ observed via APD_80_ analyses were not statistically significant **(Fig. 9I)**.

We summarized these observations of drug responsiveness by plotting the safety margin observed for each drug, using either beating prevalence (**Fig. 8J, 9J**) to obtain IC_50_ or APD_80_ prolongation (**Fig. 8K, 9K**) as the metric used to obtain EC_50_. The safety margin is defined as *in vitro* IC_50_/ETPC for prevalence (or *in vitro* EC_50_/ETPC for APD_80_), and describes the relative risk for beating abnormalities. Typically during pharmaceutical development, a safety margin of 18-20 (green line on **Fig. 8J,K** and **9J,K**) is used as a go/no-go decision for a chemical compound. None of these drugs exhibited differential pharmacology between SM and MM-pre-treated 2D monolayers in either genotype (data not shown). Safety margin analysis revealed that for WTC, culture within MPS and the subsequent maturation of MPS with fatty-acid enriched media had improved the safety margin for Verapamil (using the metric of prevalence) and Alfuzosin (using the metric of APD_80_), but no statistically significant effect on the safety margin of Flecainide (**Fig. 8J**). There was also a trend, albeit not statistically significant, toward improved safety margin of beating prevalence to Alfuzosin in MM pre-treated MPS, compared to 2D monolayers. In SCVI20, MM-treated MPS showed improved prevalence safety margins (closer to target green line) in the three drugs when compared to 2D platforms (**Fig. 9J**). APD_80_ based safety margin gave improved results for both SM and MM-treated MPS for Alfuzosin and Verapamil (although not significant for the latter one) (**Fig. 9K**). SCVI20 MM and SM treated MPS tissues were hypersensitive to Flecainide when looking at the APD_80_ safety margin, to the point that *in vitro* EC_50_ was less than the clinically reported ETPC **(Fig. 9K)**.

Besides being self-consistent with predictions of relative levels of specific currents (e.g. *I*_*Kr*_, *I*_*CaL*_) in MM vs. SM treated MPS of each genotype, these pharmacology data suggest that MPS improves the prognostic capability of hiPSC-CM, and that MM pre-treatment further augments the prognostic power of MPS. For example, although Verapamil is routinely used in the clinic, particularly for QT-interval management, it exhibits false positive toxicity in hERG-assays, and the beating prevalence of 2D monolayer cultures of hiPSC-CM are sensitized to this drug, as shown here (**Fig. 8J and 9J**) and in other studies^11,13,22^. Our data suggests that culture within MPS alone dramatically enhances the IC_50_ of this drug, eliminating the false positive toxicity seen in 2D monolayer hiPSC-CM, regardless of genotype. These observations are consistent with our previous studies^13,22^. The combination of MPS with MM gives a more accurate profile of the safe nature (thereby reducing false positive toxicity) of this drug. In contrast, the fact that Alfuzosin is sometimes observed to cause arrhythmias in the clinic^71^, suggests that the higher *in vitro* EC_50_ values observed in monolayer culture and SM-cultured MPS under-predict potential toxicity (false negative). Our findings with MM-cultured MPS, which indicate sensitization to the APD prolongation effects of Alfuzosin, suggest that MM pre-treatment enhances the ability of MPS to accurately predict the clinically observed effects of this drug.

Collectively, these data indicate that combining MPS culture with MM can reduce both false positive (Verapamil) and false negative (Alfuzosin) drug response estimates. Enhanced drug resistance is not universally observed in MPS culture, suggesting against the trivial explanation that drug availability is limiting in these 3D systems, likely due to the small and physiologically-relevant scale of our 3D microtissues ∼ 150µm in width, consistent with cardiac muscle bounded by collagen fibrils^72^. Our work suggests instead that changes in drug susceptibility are due to changes in density and function of specific ion channels that these drugs target.

## Conclusions

Despite the promise of hiPSC-derived tissue cells as genetically defined human *in vitro* models for drug development and fundamental biology, maturation of hiPSC derivatives including hiPSC-CM into adult-like cells remains an important challenge. In the present study, we demonstrated that the combination of aligned, 3D culture in MPS with fatty-acid based media synergized to promote maturation of hiPSC-CM action potential. Combining *in silico* modeling with experimental measurements provided insights into a putative mechanism linking alterations in individual ion channels and calcium handling machinery to whole-cell changes in action potential. This is the first study to induce maturation of hiPSC-CM in a tissue-chip, and, importantly, we demonstrated that maturation not only affected the baseline physiology of hiPSC-CM, but also yielded cells with pharmacology more reminiscent of adult human cardiomyocytes. Strikingly, this media had opposing effects on the action potential duration of genetically unique hiPSC-derived tissue chips, with the net result of forcing both genotypes towards an actional potential duration range that is typical for adult human ventricular cardiomyocytes. These results suggest that, as in monolayer culture of hiPSC-CM^36^, maturation with fatty-acid based media may be a prerequisite to using hiPSC-CM based tissue chips to predict how drugs will affect the adult human heart.

## Experimental Procedures

### Cell Sourcing

Two parent hiPSC were used in the majority of these studies: Wild Type C (WTC) human hiPSC harboring a single-copy of CAG-driven GCaMP6f knocked into the first Exon of the AAVS1 “safe harbor” locus^37^. The parent cell line (WTC) was reprogrammed from fibroblasts derived from a healthy 30-year-old Japanese male with a normal electrocardiogram and no known family history of heart disease and is available from the Coriell Repository (# GM25256 hiPSC from Fibroblast). The second line, Stanford University Cardiovascular Biobank Line 20 (SCVI20), was generated from a healthy, disease genotype 77-year-old Caucasian male^38^. The third cell line, SCVI273, was generated from a 43-year-old Asian female, and was used for comparison of action potential duration and calcium F/F_0_ in response to maturation media treatment. SCVI20 and SCVI273 are available from the Stanford University Cardiovascular BioBank.

### Cardiomyocyte Differentiation

hiPSC-CM were derived from pluripotent hiPSC and purified using published protocols relying on small molecular manipulation of Wnt signaling^73^, with some modifications. Briefly, frozen stocks of pluripotent cells were thawed and plated on hESC-Qualified Matrix (Corning; Corning, NY) in Essential 8 Medium (E8; Thermo Fisher, Tewksbury, MA) containing 10µM Y27632 (Peprotech; Rocky Hill, NJ). Fresh E8 without drug was added the following day. To prepare cells for differentiation, hiPSC were grown to 70-80% confluency, and then passaged three times at a constant density of 20,000 cells/cm^2^ (Burridge *et al*. 2014). During passaging, cells were singularized with Accutase (Thermo; Waltham, MA) and plated in E8 with 10µM Y27632. After pre-passaging, hiPSC were plated at a density of 25,000 cells/cm^2^, in 10µM Y27632. This was counted as “day – 3” of differentiation. At day 0, hiPSC were >90% confluent and were treated with Roswell Park Memorial Institute Medium 1640 (RPMI) containing B-27 supplement without insulin (RPMI-I), along with 8µM CHIR99021 (Peprotech) and 150µg/mL *L*-ascorbic acid (LAA). Exactly 24 hr after drug was added, medium was exchanged for RPMI-I (day 1). On day 2, medium was replaced with RPMI-I containing 5µM IWP-2 (Peprotech). On day 4, medium was exchanged for RPMI-I. RPMI containing standard B-27 supplement (RPMI-C) was added on days 6,7,9, and 11. Robust spontaneous contractile activity was typically observed on day 8 of differentiation.

On day 15 of differentiation, hiPSC-CM were singularized and cryopreserved. Briefly, cells were washed twice, for 15 minutes, with dPBS, to deplete calcium from extracellular space and sarcomeres. Next, cells were exposed to 0.25% Trypsin (Thermo Fisher) for 10-20 minutes. Cells were triturated gently at every five minutes, then pelleted (300g, 5 minutes). Cell pellets were resuspended into RPMI-C with 10µM Y27632 for counting. Cells were then pelleted a second time, and resuspended into cryopreservation medium containing 10µM Y27632, then frozen and kept in liquid nitrogen.

Two weeks before MPS experiments, hiPSC-CM were thawed and plated at a density of 100,000 cells/cm^2^ onto Matrigel, in RPMI-C with 10µM Y27632. The following day, medium was exchanged for RPMI-C. Three days after plating, monolayers were spontaneously contracting. Cells were then washed with dPBS and exposed to a cardiomyocyte selective medium depleted of glucose and pyruvate (Media-L; RPMI 1640 without glucose or sodium pyruvate, supplemented with 23mM sodium bicarbonate and 5mM Sodium *L*-lacate^74^) for a total of five days. Cells were washed with dPBS and fresh Media-L was added every other day. On the fifth day of purification, significant death of non-beating cells was observed. Cells were washed with dPBS and allowed to recover in RPMI-C for three days. Cardiomyocyte purity both before and after this procedure was characterized by flow cytometry for Cardiac Troponin T (TNNT2; **Table S1; Fig. S1**).

### Isogenic Stromal Cell Differentiation

Isogenic iPS-stromal cells (hiPSC-SC) were derived via small molecular activation of Wnt signaling in pluripotent hiPSC, followed by VEGF exposure, as described previously^75^. Briefly, hiPSC were seeded at a density of 25,000 cells/cm^2^ onto hESC-Qualified Matrigel. This was termed “day -3” of the culture. On day 0, wells were 80-100% confluent, and the medium was switched to LaSR media (Advanced F12/DMEM, 2.5mM Glutamax; 60ug/ml ascorbic acid), and 7.5uM CHIR99021 for 2 days without medium change. At day 2, the media was changed to LSR media with 50 ng/ml VEGF (Peprotech) for 2 days without medium change. On day 4, medium was replaced to LaSR media only. On day 6, cells were ready for CD31 magnetic sorting. For magnetic sorting on day 6 of differentiation, cells were rinsed with dPBS and trypsinized for 8min. Trypsin was quenched by adding EB20 media (20% FBS, 2.5mM Glutamax, KO DMEM), and cells were centrifuged (300g for 3 minutes) and re-suspended in FACS buffer (PBS, 2% FBS). CD31+ magnetic Dynabeads were added to the cell suspension at a concentration of 8 beads per CD31+ cell and left 20min on ice. The CD31 negative fraction was then expanded (maximum of ten passages) on uncoated tissue culture plastic substrates supplemented with EGM-2 media (Lonza) and characterized (**Fig. S2**). Full details on antibodies used to characterize these cells are provided in **Table S1**.

### Plating of hiPSC-CM for 2D Monolayer Studies

In 2D monolayers, hiPS-SC overgrow hiPSC-CM (data not shown). Thus, for 2D pharmacology and gene expression studies, biochemically purified hiPSC-CM were grown in monolayers. Purified cardiomyocytes were singularized with 0.25% trypsin after extensive dPBS washes. The cells were then resuspended into RPMI-C supplemented with 10µM Y27632 and plated at a density of 200,000 cells/cm^2^ onto GFR Matrigel. The following day, medium was exchanged for RPMI-C. Three days after plating, monolayers were spontaneously contracting. Cells were then exposed to either SM or MM for ten days prior to the onset of gene expression and pharmacology studies.

### Fabrication of Cardiac MPS

Microfluidic cardiac MPS systems were formed using small modifications of the protocol described in our previous work^22^ (see **Fig. 1, Fig. S6**). Briefly, two-step photolithography was used to form a chip comprised of: 1) a cell-loading port leading to a cell culture chamber with two large “anchoring posts” and several smaller micro-posts; and, 2) a media-loading port leading to media channels running alongside the cell culture chamber. The media channels and cell culture chamber (50µm high) are connected by a series of fenestrations (2µm x 2µm high/width) that provide a barrier to convective flow at defined volumetric flowrates, such that all media factors delivered to cells in the culture chamber arrive via diffusion^22^. The cardiac MPS is formed by replica molding Polydimethylsiloxane (PDMS; Sylgard 184 kit, Dow Chemical, Midland, MI) at a 10:1 ratio of Sylgard base to crosslinker from a photolithographically defined wafer. These PDMS chambers were then bonded to glass slides using oxygen plasma.

### Self-Assembly of Cardiac Microtissues within Cardiac MPS

hiPSC-CM and hiPSC-SC (passage 5 - 8) were singularized with 0.25% trypsin after extensive PBS washes. We then prepared a cocktail of 80% hiPSC-CM and 20% hiPSC-SC, at a density of 6.6×10^6^ cells/mL, in EB20 media supplemented with 10µM Y27632 and 150µg/mL *L*-ascorbic acid. 3µL of this cocktail, corresponding to 2×10^4^ cells, was injected into the cell loading inlet of each MPS. MPS were then loaded by centrifugation (300g, 3 minutes), and plugged with an SP20 steel rod to prevent cellular regurgitation from the cell chamber during media loading. Next, the same media used to resuspend cells was added to the channels of each MPS. MPS were then individually inspected, and any cell chambers that were not completely filled were filled by applying gentle pressure to the SP20 plug. This time-point was counted as MPS day 0. At MPS day 1, media was changed to RPMI with B27 supplement. At day 3 MPS were continuously fed either maturation media or standard media, using negative pressure for media exchange as described in our previous study^22^. Pump-free, gravity-driven flow was used to feed the tissues. Every 2 days, fresh media was replenished into the inlet reservoir so that hydrostatic pressure would drive a continuous flow through the media channels to the outlet reservoir. This technique is commonly used with microfluidic devices due to its simplicity and low cost (Komeya et al, 2017; ref. ^76^). Potential batch-to-batch variability in hiPSC-CM phenotype was mitigated by performing all physiology and gene expression studies with MPS and monolayers derived from a minimum of three independent differentiations.

### Robust Design Experiments to identify the composition of the optimal maturation media

We hypothesized that switching the carbon source of cardiac MPS from glucose to fatty acids could induce more mature electrophysiology of hiPSC-CM. We employed Robust Design screening to optimize four different media composition variables. Given the likelihood of these variables acting in a synergistic fashion to enhance maturation, the parametric space would require 3^4^, or 81 independent experiments (excluding the several replicates required for significant studies). To study this large space in a cost and time-effective manner within MPS, we employed Robust Design^41^ screening. With orthogonal arrays, the variable-space spanned by these 81 independent experimental conditions was explored with only 9 independent experiments. These 9 experiments were designed such that the four media input variables (levels of glucose, oleic acid, palmitic acid and BSA) were varied in an orthonormal fashion from one experiment to the next (**Table 1**). BSA (Sigma # A2153) was used directly, without being first stripped of exogenous fatty acids.

In the case where glucose was completely omitted from cardiac media, we added 10mM galactose, as previous studies have shown healthy hiPSC-CM are capable of using galactose as an ATP source^35,43^. Based on the hydrophobic nature of the primary fatty acids used as ATP sources in the heart (oleic acid and palmitic acid, respectively), we added additional BSA, above the 0.25% already contained in the B27 supplement. In all cases where fatty acids were added to media, they were first incubated at 37°C with concentrated BSA and B27 supplement to allow fatty acids to bind the extra albumin, or the 0.25% albumin already contained in B27. Beating physiology and calcium flux were assessed with high-speed microscopy as described below. Media were screened based on their ability to reduce spontaneous beat-rate, as well as the interval between peak contraction and peak relaxation during 1Hz field pacing, while maintaining a high prevalence of beating (defined as the percent of the tissue with time-averaged motion exceeding a pre-determined threshold) during pacing at 1Hz. MPS were treated with various candidate maturation media for 10 days before beating physiology was assessed.

### Image Acquisition for Beating Physiology Studies

During imaging, MPS or 2D monolayers in multi-well plates were maintained at 37°C on a stage equipped with a heating unit (Tokai Hit, Gendoji-cho, Japan). First, baseline readings of spontaneous calcium flux (GCaMP6f), and beating physiology (bright-field video) were taken. After acquiring spontaneous electrical activity, electromechanical activity under field pacing was assessed. MPS were paced via sterile, blunted stainless steel needles that were inserted into the pipette tips leading to both the media inlets and outlets. Care was taken to fill pipettes and prevent bubble formation to maintain electrical conductivity. Before recording videos, cells were paced for 10 seconds (20V, 20msec bipolar pulses at 1Hz, ION OPTIX Myopacer Field Simulator). Pacing was then maintained at the same intensity and frequency for acquiring images of MPS contracting at 1Hz.

Imaging was performed with a NIKON TE300HEM microscope equipped with a HAMAMATSU digital CMOS camera C11440 / ORCA-Flash 4.0. All videos were recorded at a framerate of 100 frames/second for a duration of 8 seconds. For GCaMP imaging, fluorescent excitation was provided by a Lumencor SpectraX Light Engine (GCaMP: Cyan LED, 470nm) and filtered with a QUAD filter (Semrock). Videos were acquired using Zeiss Zen Pro 2012 software.

### Image Analysis

Brightfield videos were analyzed for beating physiology using an updated version of our open source motion tracking software^37^ (software available at https://huebschlab.wustl.edu/resources/). The current version of the software uses tools from the open source Bioformats Toolbox^77^ to obtain image and metadata from microscopy files.

Briefly, microscopy files (Zeiss Zen, .czi) were directly read into the Matlab-based GUI, and the contractile motion of tissues was analyzed via an exhaustive-search block-matching optical flow algorithm that compared the position of 8×8 pixel macroblocks at frame *i* to their position at frame *i*+5 (corresponding to the motion in 50msec). Motion vectors were used to calculate beat-rate, beating interval (defined as the time delay between maximum contraction velocity and maximum relaxation velocity, which is directly proportional to action potential duration), and beating prevalence. Beating prevalence was defined as the percentage of macroblocks within a region-of-interest (ROI) with a time-averaged contraction speed that exceeds a defined threshold (2µm/sec was defined empirically as a universal threshold for all MPS analyzed). ROI were selected to include the entire cell culture chamber of the MPS.

GCaMP data were quantified using in-house Matlab code that was developed based on previous work by Laughner and colleagues^54,78^. GCaMP videos were analyzed for τ_75_ decay time (time for calcium amplitude to go from maximum to 25% of maximum), Full Width Half Maximum (FWHM, time for calcium amplitude to go from 50% of maximum during upstroke, to 50% of maximum during decay) as well as peak intensity, a metric of total calcium influx. For spontaneously beating cells, data on beating interval and calcium transient decay times were rate corrected using Fridericia’s method^79^.

### Optical Measurement of Action Potentials

BeRST-1 dye was synthesized, and purity verified, as previously described^80^. For action potential recording, MPS were first labeled overnight with 2.5µM BeRST-1. The following day, MPS were equilibrated to media without dye before imaging (RPMI-C without phenol red) as described above, using a Red LED (640nm). For monolayer experiments, cells were labeled with 500nM BeRST-1 for 1h at 37°C, and then equilibrated to RPMI-C without phenol red. After acquiring videos of spontaneous and 1Hz paced activity at 100 Hz for 8 seconds, BeRST-1 videos were analyzed using similar Matlab code as was used for GCaMP analysis^78^. BeRST-1 videos were analyzed for 80% Action Potential Duration (APD_80_). Reported values of APD_80_ (**Fig. 3**) are for MPS or monolayers paced at 1Hz.

### MPS Tissue Isolation and Immunofluorescence Imaging

Tissues were treated with SM or MM for 10 days. On day 10, MPS were flushed for 10 minutes with PBS at 25°C. Following this wash, 4% PFA was added to the media channel for 15min to fix the tissues. MPS were washed with PBS twice for 5min after that and were then carefully cut with a clean scalpel, to open the device and expose the tissue. At this point, the PDMS component had the tissue structure attached to it. The tissues were then stained by submerging PDMS blocks in different staining solutions: First, tissues were blocked with blocking buffer (1% BSA 10% FBS 0.5% Triton 0.05% sodium azide) overnight at 4°C. The next day, they were submerged the primary antibodies (Mouse anti α-actinin, Life technologies 41811; Rabbit anti-GAPDH, abcam 9485; Mouse anti-mitochondria, abcam 92824; Mouse anti β-myosin heavy chain, abcam 97715; Rabbit anti myosin Light Chain 2V (MLC2V), Proteintech 10906-1-AP) 1:100 concentration in blocking buffer) for 48h at 4°C. Tissues were then washed twice at 25°C in blocking buffer for 2-3 hours and washed a third time at 4°C overnight. The secondary antibodies (Goat anti-mouse IgG Alexa 568 H+L, Life Technology a11004; Goat anti-rabbit IgG Alexa 488 H+L, Life Technology a11008) along with 1:600 DRAQ5 (Abcam, ab108410) was incubated in blocking buffer for 24h. Tissues were then washed twice at 25°C in blocking buffer for 2-3 hours and a third time at 4°C overnight before tissues were imaged.

Both WTC and SCVI20 tissues were imaged with Opera Phenix™ High Content Screening System. All images were taken through proprietary Synchrony™ Optics 63x water immersion lens. Images were acquired using Harmony software. We imaged both DRAQ5 and α-actinin for sarcomere alignment using 640nm and 546nm lasers respectively. We performed z-stacks over 60μm with a step-size of 0.5μm for α-actinin or GAPDH stains. Mitotracker and anti-mitochondrial antibody were imaged with a step size of 0.3μm. Post imaging processing was performed on ImageJ to enhance contrast and decrease background fluorescence. The same post-processing was performed for SM and MM tissues to allow direct comparison between them.

To analyze the regularity of sarcomeres from staining of sarcomeric α-Actinin, we applied Fast Fourier-Transform (FFT) based methods^54^ to cellular regions of the MPS that had a constant size (100 x 100 pixels). Next, the real component of the FFT was smoothed with a 3×3 Gaussian filter, and the mean intensity was calculated as a function of radial distance from the center of the centered real-component of the FFT. Structures with regularly repeating features (e.g. sarcomeres) produce distinct bands when analyzed in this manner, resulting in local increases in intensity at specific radial distance. These local intensity increases were quantified^43,54^. Code is available from the authors upon request. Representative fields of interest for intensity quantification and analysis of sarcomere regularity (**Fig. S5**) were selected and analyzed by a user blinded to the experimental condition.

### Analysis of Mitochondrial Morphology and Potential

Mitochondrial potential was analyzed in 2D monolayers and MPS using MitoTracker Red (M7512 Thermo Scientific). Live MPS and monolayer were incubated with Mitotracker for 30min at 37C in RPMI 1640 supplement with insulin. Samples were then washed with PBS for 10min before being fixed in 4% PFA for 15min and washed again with PBS. Monolayers were directly imaged after that and tissues were isolated from the MPS as described above and placed into wells of 24-well plates with PBS until they were imaged.

### Measurement of contraction force

Micro-molded PDMS pillars were added to the cell chambers, so that the tissue would deflect them upon each contraction (**Fig. 1B, S6A,B**). By considering each pillar as a cantilever beam fixed at one end and uniformly loaded with horizontal forces along its height, one can apply the Euler-Bernouilli formula for uniformly distributed load and deduce the contraction force from the pillar’s elastic modulus, deflection and dimensions. For WTC-MPS, pillar deflection was calculated with ImageJ by measuring the distance between the pillar’s centroid coordinates before and after contraction. Same scale (0.5 pixel/micron) was used for all measurements.

To facilitate higher-throughput measurements, a new python-based software was developed to automatically identify pillars in the tissue. The software combined information derived from circle Hough transform^81^ and custom template matching^82^ algorithms (**Fig. S6C**) to detect the initial coordinates of the pillars at frame 0 of each recording. These initial positions were then fed to an enhanced version of our original tracking algorithm^37^ to retrieve the absolute motion of the pillars during contractions (**Fig. S6D**). The motion was then converted to contraction force by using Euler’s beam theory as described previously. This higher throughput software was used for SCVI20 MPS force analysis.

### Pharmacology in MPS and 2D Monolayers

To avoid any possible confounding effects that different albumin or lipid content might have on drug bioavailability^83^, for all pharmacology, MPS were first equilibrated to phenol red free RPMI with B-27 (SM) containing vehicle control (DMSO, methanol, or water, to a final concentration of 0.1% v/v). On the day upon which studies were performed, freshly measured drug was dissolved into DMSO, except for Flecainide, which came as a methanol stock solution, and Verapamil or Isoproterenol, which were dissolved directly into media. For testing inotropic responsiveness to extracellular calcium concentration, Tyrode’s saline (0.1g/L anhydrous MgCl_2_, 0.2g/L KCl, 8g/L NaCl, 50mg/L anhydrous monobasic sodium phosphase, 1g/L D-glucose) was used in lieu of RPMI-C, as in previous studies of inotropic responsiveness of macroscale and miniaturized EHM^13,17^. After recording activity in zero-dose vehicle condition, media were exchanged for the lowest drug dose, and MPS were incubated for 30 minutes at 37°C. Spontaneous activity, and activity with 1 Hz pacing, were recorded as described above. This was repeated for each dose escalation of drug. Drug dose was escalated until all spontaneous and paced activity ceased, or a dose of 10µM was reached.

Media on monolayers of hiPSC-CM was replaced with phenol red free RPMI-C containing 1µM BeRST-1, and monolayers were incubated for 30 minutes at 37°C (5% CO_2_). Medium was next replaced with RPMI-C supplemented with the vehicle used to dissolve drug (water, methanol or DMSO, to a final concentration of no more than 0.1% v/v). Similar to dose-escalation studies in MPS, new drug was added and allowed to equilibrate to each increasing drug dose over 30 minutes intervals at 37°C, 5% CO_2_. After equilibrating monolayers to vehicle and to each dose of drug, spontaneous beating physiology, calcium flux and action potentials were collected in bright-field, GCaMP and BeRST-1 channels, respectively. Next, cells were paced at 1Hz to collect these same three parameters.

### Gene Expression in Monolayer Culture

To characterize gene expression during hiPSC-SC differentiation, cells at various stages of differentiation of hiPSC to endothelial cells were trypsinized, pelleted and lysed (Qiagen RLT lysis buffer supplemented with 1% β-Mercaptoethanol. RNA was recovered using spin columns (Qiagen MicroRNAeasy® kit), with on-column DNAse I digest performed according to the manufacturer’s protocol. Purified hiPSC-CM were plated to a density of 200,000 cells/cm^2^ on Matrigel coated plates in RPMI-C containing 10µM Y27632. One day after plating medium was exchanged for RPMI-C. Two days following this, monolayers had recovered spontaneous beating, and cells were treated for ten days with either Standard Media (SM) or Maturation Media (MM). Media was exchanged every 3 days. On day 10, cells were washed with PBS and RNA was recovered in the same manner as for monolayer hiPSC-SC. Following RNA recovery from 2D cultures, 500ng of recovered RNA was used to produce cDNA using the SuperScript III kit with Oligo-dT primers (Life Technologies). The obtained cDNA was used to perform SYBR Green and Taqman real-time PCR analysis with the probes described in **Supplemental Tables S1 and S2**. Commercial polyA-RNA obtained from 15 pooled male/female Caucasians adult human left ventricle (Clonetech, Mountain View, CA) was used as a positive control for expression of cardiomyocyte ion channels, as described previously by Liang^11^.

### Gene Expression in MPS

Purified hiPSC-CM were combined with expanded hiPSC-SC as described above for initial optimization studies and cultured for ten days in either standard SM or MM. We first optimized protocols for isolating high-quality (R.I.N. > 8.5) RNA from MPS (data not shown). RNA was extracted from tissue using methods similar to those previously applied for macroscale engineered heart muscle preparations^18^. Briefly, on day 10, MPS were flushed for 10 minutes with PBS at 25°C. Following this wash, MPS were carefully cut with a sterile scalpel, to separate the cell culture chamber from the cell loading chamber of the MPS, and to open the device. The PDMS component, with microtissue attached, was transferred to a microcentrifuge tube, and pooled with up to six other microtissues in lysis buffer from the RNAqueous kit (Thermo). Immediately after adding the tissues to lysis buffer, the microcentrifuge tube containing them was flash frozen in liquid nitrogen. Next, RNA was retrieved from samples by following manufacturer instructions on the RNAqueous kit, followed by DNAse I digestion (Ambion). The yield and quality of RNA were assessed with Qbit and Bioanalyzer, and with optimized methods, we routinely achieved RNA Integrity Numbers above 9. RNA isolated from MPS was amplified using a SMARTeR stranded Pico Input Total RNA library prep kit (Clonetech). The cDNA library products were then diluted by a factor of 10 into sterile water for direct quantitative PCR analysis of relative gene expression. RNA from positive controls was reverse-transcribed and amplified using the same kit. For qPCR analysis, cDNA libraries were diluted by a factor of ten into RNA grade water so that gene expression would fall within the linear assay range.

### Mathematical Modeling

Time-series of AP and Ca^2+^ flux from MPS paced at 1Hz were inverted to a mathematical model of ion channel activity and calcium dynamics to obtain simulated estimates of channel conductance and calcium handling as described in our recent publication^64^. Briefly, a modified version of a model of an immature stem cell^7^ was used to calculate the predicted voltage and calcium dynamics. Parameters of this model, specifically maximal channel conductances, intracellular calcium diffusion terms, and surface to area ratios, were then iteratively perturbed until the error between the measured waveforms and simulated waveforms was minimized. Resulting parameters and produced action potential models were then plotted by group to provide an explanation for mechanistic reasons for changes in action potential. It is known that parameterization of such models based on MPS data is challenging and may yield non-unique results; different channel conductances may results in similar voltage output of the model. This issue is discussed by Jæger in a previous study (Jæger *et al. Chaos* 2019; ref. ^84^) where a method for identifying undetectable parameters was derived. In the present work we have used an inversion with regularization^85^ to handle non-unique solutions.

### Statistical Analysis

Direct comparisons were made by unpaired student’s *t*-test, with Holm-Bonferonni correction for multiple comparisons. All curve fitting was done using GraphPad Prism. IC_50_ and EC_50_ curves were fit to four-parameter models. When these models yielded ambiguous fits (**Fig. 5C,D,F,G**), a three-parameter model was used. To analyze the currents and calcium flux parameters generated from model fits of action potential and calcium flux waveforms (Fig. 6), we performed a global 1-way ANOVA analysis followed by post-hoc Tukey tests. Gene expression data were statistically analyzed with ClustVis (web tool for clustering multivariate data) and GraphPad Prism. Overall PCR data were plotted on ClustVis to obtain heatmaps of the gene expression for maturation media treated MPS relative to standard media values. The genes within 90% percentile of differential expression were then selected and plotted on GraphPad Prism where an unpaired non-parametric t-test was performed to compare standard media and maturation media values using the Holm-Sidak method. Significance was determined with *p*-value < 0.05.

## Acknowledgements

This work was funded in part by the California Institute for Regenerative Medicine DISC2-10090 (K.E.H.), NIH-NHLBI HL130417 (K.E.H.), NIH-NIGMS R35GM1195855 (E.W.M.), NIH-NIGMS T32GM066698 (S.B.), the Research Council of Norway INTPART Project 249885, the SUURPh program funded by the Norwegian Ministry of Education and Research, and the Peder Sather Center for Advanced Study (UC Berkeley). We thank Mary West (UC Berkeley) for assistance with image analysis and flow cytometry and Silvio Weber (Technische Universität Dresden) and Stacey Renschler (Washington University in St. Louis) for helpful advice on RNA isolation, cDNA amplification and data analysis. We thank Yoram Rudy, Jon Silva and Jianmin Cui (Washington University in St. Louis) for critical discussion on action potential acquisition, mathematical modeling and data analysis. We thank Bruce Conklin (Gladstone Institutes, San Francisco, USA) for technical advice on the WTC iPSC line, and Joseph Wu (Stanford University) and the Stanford University Cardiovascular BioBank for providing the SCVI20 iPSC line and providing technical advice regarding this line.

Professors Kevin E. Healy, Sam Wall, Andrew Edwards and Nathaniel Huebsch have a financial relationship with Organos Inc. and both he and the company may benefit from commercialization of the results of this research.

