## Supplemental Materials for "Metabolically-Driven Maturation of hiPSC-Cell Derived Cardiac Chip"

**Supporting Materials**


Supplemental Methods

Table S1. Antibodies

Table S2. Primer Pairs for SYBR Green Based Gene Expression Analysis

Table S3. Probes used for Taqman Based Gene Expression Analysis

Figure S1. Characterization of the Purity of iPSC-CM Obtained by Lactate Treatment of Cryopreserved iPSC-Cardiomyocyte Differentiation

Figure S2. Derivation of hiPSC Derived Cardiomyocytes and Isogenic Stromal Cells

Figure S3. Additional characterization of action potentials in MPS and Monolayers

Figure S4. Metabolic response of iPSC cardiomyocyte monolayers to Maturation Media

Figure S5. Quantitative analysis of sarcomere regularity and intensity of expression of Sarcomere proteins associated with mature contractile machinery in MPS

Figure S6. Analysis of Force Developed by Cardiac Microphysiological Systems

Figure S7. Quantification of GAPDH protein levels in MPS

Figure S8. Experimental Analysis of Steady State Sarcoplasmic Reticulum Calcium Levels in MPS treated with Maturation Media

### Supplemental Methods

#### Formation of Microphysiological Systems Using Defined hiPSC Derived Cardiomyocytes and Stromal Cells

In previous studies, we used cardiomyocytes derived from Wnt-mediated hiPSC cardiac specification[1], along with byproduct stromal cells, with no purification steps. Here, tissue formulation was refined by combining biochemically purified hiPSC-CM with isogenic stromal cells (hiPSC-SC). Methods were first optimized to determine the most reproducible procedure for obtaining a high yield of purified hiPSC-CM. We found that cryopreserving hiPSC-CM at day 15 of differentiation, and then treating cells with glucose depleted, lactate enriched media after thawing was the most reliable method. After lactate purification, hiPSC-CM stayed pure for more than one week (**Fig. S1**).

To obtain a defined hiPSC-SC population, we used the stromal biproducts of endothelial lineage specification of isogenic hiPSC. These hiPSC-derived stromal cells (hiPSC-SC) have a similar lineage to cardiomyocytes, sharing a common MESP1 positive mesodermal progenitor (**Fig. S2A,B**). After expansion, hiPSC-SC exhibited significant transcription of IGF and BMP-2 (**Fig. S2B**), two cytokines known to be important for cardiac development[2,3].

Immunofluorescence analysis suggested robust expression, with little observable heterogeneity, of the stromal markers α-Smooth Muscle Actin (αSMA), SM22, CD90 and Vimentin. Importantly, hiPSC-SC also express N-Cadherin and the gap junction marker Connexin 43 (Cx43/GJA1; **Fig. S2C**), suggesting these cells would be capable of coupling effectively to cardiomyocytes in micro-tissues. Finally, hiPSC-SC produced ECM transcripts including collagen (I & IV), laminin and fibronectin, although the level of ECM deposition appeared to be substantially higher for collagen IV as compared to collagen I (**Fig. S2C**). These tissue-specific ECM proteins are reminiscent of the composition of ECM in the developing heart[4].

Using these purified, defined cell populations, we fabricated MPS that mimicked the mass composition of the human heart by combining 80% hiPSC-CM and 20% hiPSC-SC. These tissues exhibited a high level of cellular alignment and sarcomere organization (**Fig. S2D**).

#### Analysis of Force Developed by Cardiac Microphysiological Systems

To facilitate direct measurement of cardiomyocyte contractile forces, micro-molded polydimethylsiloxane (PDMS) pillars were added to the cell chambers, so that the tissue would deflect them upon each contraction (**Fig. S6A**).

The presence and size of pillars was verified by Scanning Electron Microscopy (SEM). Prior to SEM imaging, the PDMS MPS devices were coated with a thin layer of gold-palladium (AuPd) to make the surface conductive. A Cressington 108 SEM Sputter Coater was used for it. A 0.06 mbar Argon atmosphere was created and the samples were sputtered for 60s at 10mA (1kV) at a distance of 4cm from the target, resulting in 100Å AuPd. Samples were then mounted on a SEM holder using copper tape and adhesive carbon disks. SEM images were taken on a FEI Quanta 3D FEG Dual beam (SEM/FIB) system in standard high vaccum mode with 20kV/120pA electron beam. The imaging parameters for our PDMS MPS were the following: 350x magnification, 45 degrees tilt, working distance 9.5mm.

When micro-tissues were formed within micro-pillar modified MPS, the twitch force of cellular contraction deflected the pillars. By considering each pillar as a cantilever beam fixed at one end and uniformly loaded with horizontal forces along its height, one can apply the Euler-Bernouilli formula for uniformly distributed load and deduce the contraction force from the pillar's elastic modulus, deflection and dimensions (**Fig. S6B**). Pillar deflection was calculated in ImageJ by measuring the distance between the pillar's centroid coordinates at zero and at maximal contraction.

For automated pillar tracking, Python-based algorithms were developed. Pillar deflection is automatically calculated in a two-step process: first, initial pillar position is detected (**Fig. S6C**). The algorithm takes as input the raw bright-field image. By combining the information of the circle Hough transform algorithm to find circular shapes in the input and a template matching algorithm using a scaled and rotated binary template of the chip design and the circle, it is possible to generate a correlation map predicting the initial position of the pillars. Second, pillar deflection is computed automatically (**Fig. S6D**). The tracking software makes it possible to follow the displacement of the pillar in time by taking the mean motion of a population of 200 tracking points placed all over the pillar before contraction and during Supra-max of the contraction with tracking points distributed on the pillar.

**Staining of Nascent T-Tubules in Cardiac MPS**

MPS were rinsed three times for five minutes with PBS, and then fixed with 4% paraformaldehyde (PFA) for 15 minutes. After 3 more five-minute PBS washes, MPS were carefully cut open with a scalpel and transferred to multiwell plates. After one additional PBS wash, MPS were stained with 20μg/mL Alexa Fluor 488 conjugated Wheat Germ Agglutinin (WGA-488; Thermo Scientific) for 30 minutes. After 3 additional washes in PBS, MPS were mounted on glass slides with ProLong Gold Antifade reagent (Thermo Scientific).

Imaging was performed on an Olympus FluoView 1000 confocal microscope (Olympus, Center Valley, PA). The sample was laser-illuminated at 488 nm and fluorescent emission (525 +/- 25 nm) was collected via 60x/1.42 NA oil immersion objective (0.414 um/pixel, 100 nm confocal pinhole). Images were collected in 2D mode at lower laser powers (0.5-3%) and long pixel dwell times (40-100 us) to optimize contrast in these fixed samples.

**Supplemental Figure Legends**

**Supplemental Figure 1. Characterization of the Purity of iPSC-CM Obtained by Lactate Treatment of Cryopreserved iPSC-Cardiomyocyte Differentiation. A)** Representative plots indicating how gating was performing to identify cells from FSC and SSC plots (left), and contour plots of negative control (center) and lactate-purified cardiomyocytes (right). Because **B)** Representative histogram plots of the relative number of Cardiac Troponin (TNNT2) positive cells present immediately after lactate treatment (left), six days (center) and 180 days (right). Only after extremely long time-periods is there a noticeable reduction in cell purity. All MPS were formed with iPSC-CM within six days of lactate treatment. **C)** Time course of purity of three different batches of iPSC-CM thawed immediately after differentiation and subsequently purified with lactate treatment. **D)** Representative immunofluorescence micrograph of an iPSC-CM after lactate purification. Image on right is magnified to display sarcomeres stained with antibodies against Sarcomeric α-Actinin (green). Error bars: *SD*, $n = 3$.

**Supplemental Figure 2. Development and Characterization of iPSC-Stromal Cells**. **A)** Differentiation tree depicting the lineage of iPSC-cardiomyocytes (hiPS-CM) and the isogenic hiPSC-derived stromal cell population (hiPSC-SC). Specific biomarkers (blue) were verified by qRT-PCR. **B)** Brewer plot identifying gene expression patterns over the course of differentiation of hiPSC into hiPSC-SC, with hiPSC-CM included on the plot for comparison. **C)** Immunofluorescence molecular characterization of hiPSC-SC. These hiPSC-SC were markedly positive for all stromal markers shown, while markers of smooth muscle (Calponin) and endothelial cells (CD31) were not detected. iPS-SC also produce key ECM proteins: Laminin, Fibronectin and Collagen IV, while substantial Collagen I was not detected. **D)** Representative fluorescence micrograph of cardiac MPS, which shows highly organized, aligned cardiomyocytes stained for sarcomeric α-Actinin (ACTN2, green). Cells were counterstained for nuclei with DAPI (blue) in all fluorescence micrographs. Scale bars: **C**) 500μm **D**) left panel, 20μm and right panel, 10μm.

**Supplemental Figure 3. Additional information about the media screen and representative 2D traces of MM-treated monolayers. A-B)** Quantification of action potential time from 20% above baseline to peak ($Upstroke_{80}$) in **A)** WTC MPS and 2D monolayers and **B)** SCVI20 MPS. **C,D** Quantification of C) beat-rate corrected $cAPD_{80}$ and D) background corrected calcium amplitude ($F/F_0$) in MPS formed from SCVI273 iPSC-CM and stromal cells. **E,F)** Representative voltage tracings for 2D monolayers of **E)** WTC and **F)** SCVI20 cell lines cultured for one week in Maturation Media (MM). Voltage tracings were obtained by overnight labeling of monolayers with BeRST-1. **G-H)** Analysis of changes in **G)** APD80 and **H)** contractile prevalence in MM-pretreated MPS that resulted from modulating the levels of palmitate and albumin in MM. Removing both palmitate and albumin from MM (M1) resulted in MPS with APD80 that were significantly higher than APD80 of MM-treated MPS, and which were no different from APD80 of SM-treated MPS. Removal of Palmitate alone (M2), or of albumin alone (M3) led to a new medium that exhibited APD80 significantly less than MM ($p < 0.05$). Compared to MM, M2 treated MPS exhibited slightly reduced beating prevalence, whereas this metric was enhanced for M3 treated MPS, although these changes were not statistically significant. Slight reduction of the albumin content of MM from 2.5% to 1% (M4) did not have significant effects on APD80 or prevalence of motion in MPS, compared to those treated with MM. All data: plot of all points with median, $n > 5$. (** $p < 0.01$, 2-way t-test with Holm-Bonferonni correction for multiple comparisons).

**Supplemental Figure 4. Metabolic analysis of MM and SM-treated 2D iPSC-CM Monolayers. A)** Representative Oxygen Consumption Rate (OCR) tracings of SCVI20 iPSC-CM monolayers after culture in MM (blue) or SM (red) for 10 days. **B,C)** Quantification of reserve OCR (change in OCR from baseline with FCCP treatment) and total ATP capacity indicate a shift toward ß-oxidation with MM in 2D monolayers. **D-G)** Representative images of 2D **D,E)** WTC and **F,G)** SCVI20 iPSC-CM monolayers stained with MitoTrackerRed (left; red) and anti-mitochondrial antibodies (right; green).

**Supplemental Figure 5. Expression and localization of sarcomere proteins in MM-treated MPS. A-B)** Fourier domain-based quantification of sarcomeric order in MPS treated with SM versus MM for (**A**) WTC or **(B)** SCVI20 cell line treated with MM (blue) or SM (red) for 10 days. **C,D)** Quantification of protein expression (antibody staining with analysis by a condition-blinded user) for **C)** MYH7 and **D)** MLC-2v in MPS treated for ten days with MM (blue) or SM (red). **E-G)** Representative micrographs of **E)** WTC and **F,G)** SCVI20 MPS after staining for **E,F)** MYH7 and **G)** MLC-2v. Antibody staining in green, with blue Draq5 nuclear counterstain. Scale bars: 20μm.

**Supplemental Figure 6. Characterizing Force Developed by Contracting Cardiac Microphysiological Systems. A)** Illustration of the micro-molded PDMS pillars in the heart-on-chip platform. Left to right shows (i) one empty chamber imaged via brightfield microscopy, (ii) SEM image of the magnified pillar array. **B)** Pillar deformation can be described by the formula for uniformly distributed load on a cantilever beam, where $\delta max$ [m], the maximal deflection is proportional to the line pressure load q [N/m], the length of the beam L[m] and inversely proportional to the Young's modulus E [N/m$^2$] (2.63 [Mpa] for 1:10 PDMS network) and the second moment of area I [m$^4$], for a cylinder of radius R [m]. From this formula, it is possible to isolate final contraction force F [N]. **C-D)** Automated pillar tracking and deflection analysis to measure tissue contractile force. **C) Initial Pillar Position detection**. Left to right (i) The algorithm takes as input the raw bright-field image on which (ii) Hough circle transform (HCT) algorithm is run. This algorithm is able to detect circles in raw images by applying an edge detection filter. Each edge point becomes a center for a new circle. The intersection of all these new circles will give the central position of the initial circular shape in the raw image. (iii) A template matching algorithm is then applied to the raw image with HCT detected circles. The template given is a binary pillar pattern template that will be rotated, zoomed and scaled to find the highest matching probability with HCT circles. The latter will generate a correlation map (iv) predicting the most likely initial position of the pillars (v). **D) Automated pillar tracking**: The tracking software assess pillar displacement by measuring mean motion of a population of 200 tracking points placed all over the pillar before contraction (left) and during maximum of the contraction with tracking points distributed on the pillar (right). Scale bars: A) middle panel, 50µm and right panel, 50µm.

**Supplemental Figure 7. Analysis of Glyceraldehyde 3-phosphate dehydrogenase (GAPDH) Expression in MM treated MPS. A)** Quantification of GAPDH protein expression (immunostaining) as (left) raw fluorescence intensity, and (right) pixel-wise ratio of intensity of GAPDH / ACTN2. **B-E)** Representative micrographs of MPS stained for GAPDH (green) with Draq5 nuclear counterstain (blue) for **B,C)** WTC and **D,E)** SCVI20 MPS. Scale bars: 20μm.

**Supplemental Figure 8. Effects of ryanodine on MM and SM pre-treated MPS. A-D)** WTC MPS pre-treated with MM and SM were treated with 10μM ryanodine, and **A)** Normalized beat rate, **B)** Normalized $F/F_0$, **C)** Normalized beat-rate corrected decay time ($\tau_{75}$) and **D)** upstroke duration were quantified for spontaneous calcium transients (GCaMP6f).

**Supplemental Figure 9. Effects of thapsigargin on MM and SM pre-treated MPS. A-D)** WTC MPS pre-treated with MM and SM were treated with 10μM thapsigargin, and **A)** Normalized beat rate, **B)** Normalized $F/F_0$, **C)** Normalized beat-rate corrected decay time ($\tau_{75}$) and **D)** upstroke duration were quantified for spontaneous calcium transients (GCaMP6f).

**Supplemental Figure 10. Qualitative analysis of nascent T-tubules in MM and SM pre-treated MPS.** Cardiac MPS were stained *in situ* with 20μg/mL Alexa Fluor 488 labeld Wheat Germ Agglutinin (WGA-488) with Draq5 nuclear counterstain (blue) to probe for potential T-tubules. **A-C)** Representative staining of MM treated SCVI20 MPS showed some internal membrane invaginations (between the border of the cells and the nucleus, red arrows). **D,E)** Representative staining of SM treated SCVI20 MPS showed less prominent staining of internal membrane invaginations. Scale bar: 5μm.

**Supplemental Figure 11. Calcium Flux Amplitude and Kinetics Changes in Flecainide Treated SCVI20 Microphysiological Systems. A-C)** Analysis of calcium transient amplitude and kinetics in SCVI20 MPS pre-treated with either standard media (SM) or maturation media (MM) and then exposed to either 100nM or 1μM flecainide. Calcium transients were analyzed for **A)** upstroke time**, B)** $\tau_{75}$, the time required for calcium amplitude to decay from the maximum value to 30% of the maximum value, and **C)** maximum amplitude of the calcium transient. Values are normalized to the timing (upstroke time and $\tau_{75}$) and amplitude of the calcium transient for the same MPS treated with vehicle. **D-E)** show representative traces of SM-treated **(D)** or MM-treated **(E)** tissues exposed to 100nM flecainide. SM shows clear instances of delayed-after depolarizations (DADs), whereas MM did not show any.

**Table S1. Antibodies**

| Antibody Target | Species Raised in | Vendor | Cat. No. |
|---|---|---|---|
| Sarcomeric α-actinin | Mouse (clone EA-53) | Sigma Aldrich | A7811 |
| Laminin | Rabbit (polyclonal) | Abcam | ab14055 |
| Cardiac troponin T (FC) | Mouse (Clone 13-11) | Thermo Scientific | MS295P |
| Connexin 43 | Rabbit | Sigma Aldrich | C6219 |
| SM22 | Rabbit | Abcam | Ab14106 |
| Vimentin | Mouse | Zymed | 08-0052 |
| α-Smooth Muscle Actin (αSMA) | Rabbit | Abcam | Ab5694 |
| CD90 | Mouse (Clone 5E10) | eBioscience | 14-090-80 |
| N-Cadherin | Rabbit (polyclonal) | Abcam | ab23751 |
| Fibronectin (FN) | Rabbit (polyclonal) | Abcam | ab23751 |
| Collagen IV (Col IV) | Mouse (Clone 23IIC3) | Millipore | MAB1910 |
| Collagen I (Col I) | Rabbit (polyclonal) | Abcam | ab21286 |
| Calponin | Rabbit (polyclonal) | Abcam | ab46794 |
| CD31 | Mouse (Clone M89D3) | BD Biosciences | 558068 |
| Myosin Heavy Chain (MYH7) | Mouse (Clone M14) | Abcam | ab97715 |
| Myosin Light Chain, Ventricular Isoform (MYL2) | Rabbit (polyclonal) | ProteinTech | 10906-1-AP |

**Table S2. Primer Pairs used for SYBR Green Based Gene Expression**

| Gene | Forward Primer | Reverse Primer |
|---|---|---|
| **VIM** | GACGCCATCAACACCGAGTT | CTTTGTCGTTGGTTAGCTGGT |
| **BMP2** | TAATTCGGTGATGGAAACTG | CCCAGAAGGAAGTACATTTG |
| **IGF1** | CCCAGAAGGAAGTACATTTG | GTTTAACAGGTAACTCGTGC |
| **COLA1** | GCTATGATGAGAAATCAACCG | TCATCTCCATTCTTTCCAGG |
| **COL4A1** | AAAGGGAGATCAAGGGATAG | TCACCTTTTTCTCCAGGTAG |
| **GATA2** | CTACTAAAGCTGCACAATG | CTTTCTTGCTCTTCTTGGAC |

All other SYBR green primer pairs were provided as part of a custom gene array (Super Array Biosciences)

**Table S3. Taqman Probes used for Gene Expression**

| Gene | Taqman Probe ID (Thermo Scientific) |
|---|---|
| SCN5A | Hs00165693_m1 |
| KCNJ2 | Hs00265315_m1 |
| KCNJ3 | Hs00158421_m1 |
| CACNA1C | Hs00167681_m1 |
| HCN2 | Hs00606903_m1 |
| HCN4 | Hs00175760_m1 |
| KCNIP2 | Hs01552688_g1 |
| KCND3 | Hs00986860_m1 |
| KCNQ1 | Hs00923522_m1 |
| KCNA5 | Hs00969279_s1 |
| TNNT2 | Hs00943911_m1 |
| MYH7 | Hs01110632_m1 |
| RYR2 | Hs00181461_m1 |
| MYH6 | Hs01101425_m1 |
| SLN | Hs00161903_m1 |
| CACNB2 | Hs01100744_m1 |
| LDHA | Hs01378790_g1 |
| ADHFE1 | Hs00329084_m1 |
| ENO1 | Hs00361415_m1 |
| HEY2 | Hs01012057_m1 |
| PECAM1 | Hs01065279_m1 |
| SERCA (ATP2A2) | Hs00544877_m1 |
| CACNB1 | Hs00609503_g1 |
| NPPB | Hs00173590_m1 |
| NR2F2 | Hs00819630_m1 |
| KCNH2 | Hs04234270_g1 |
| COX41 | Hs00971639_m1 |
| MYL7 | Hs01085598_g1 |
| MYL2 | Hs00166405_m1 |
| KCND2 | Hs01054873_m1 |
| ALDH1A1 | Hs00946916_m1 |
| GAPDH | Hs02758991_g1 |

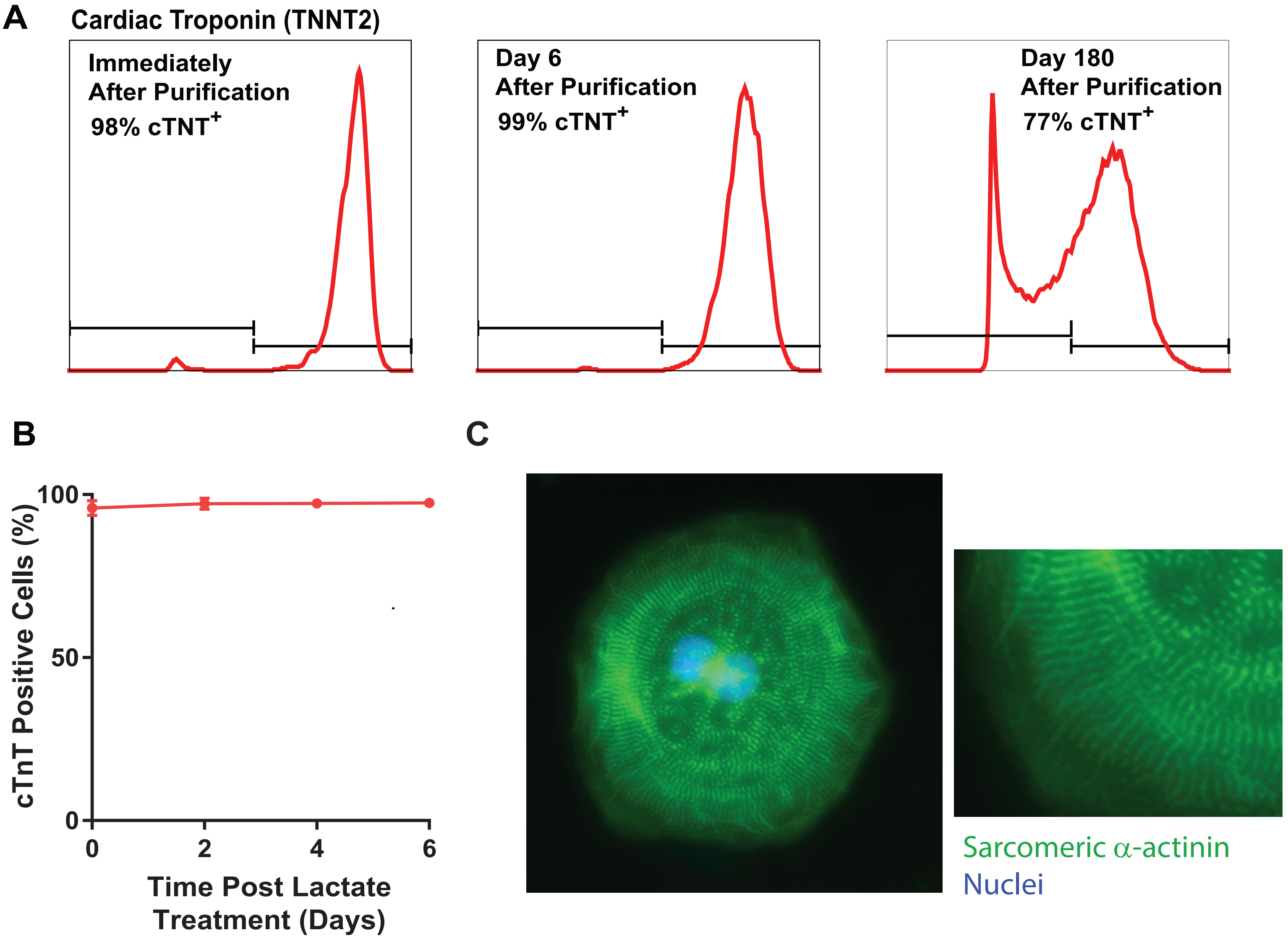


Supp. Figure 1

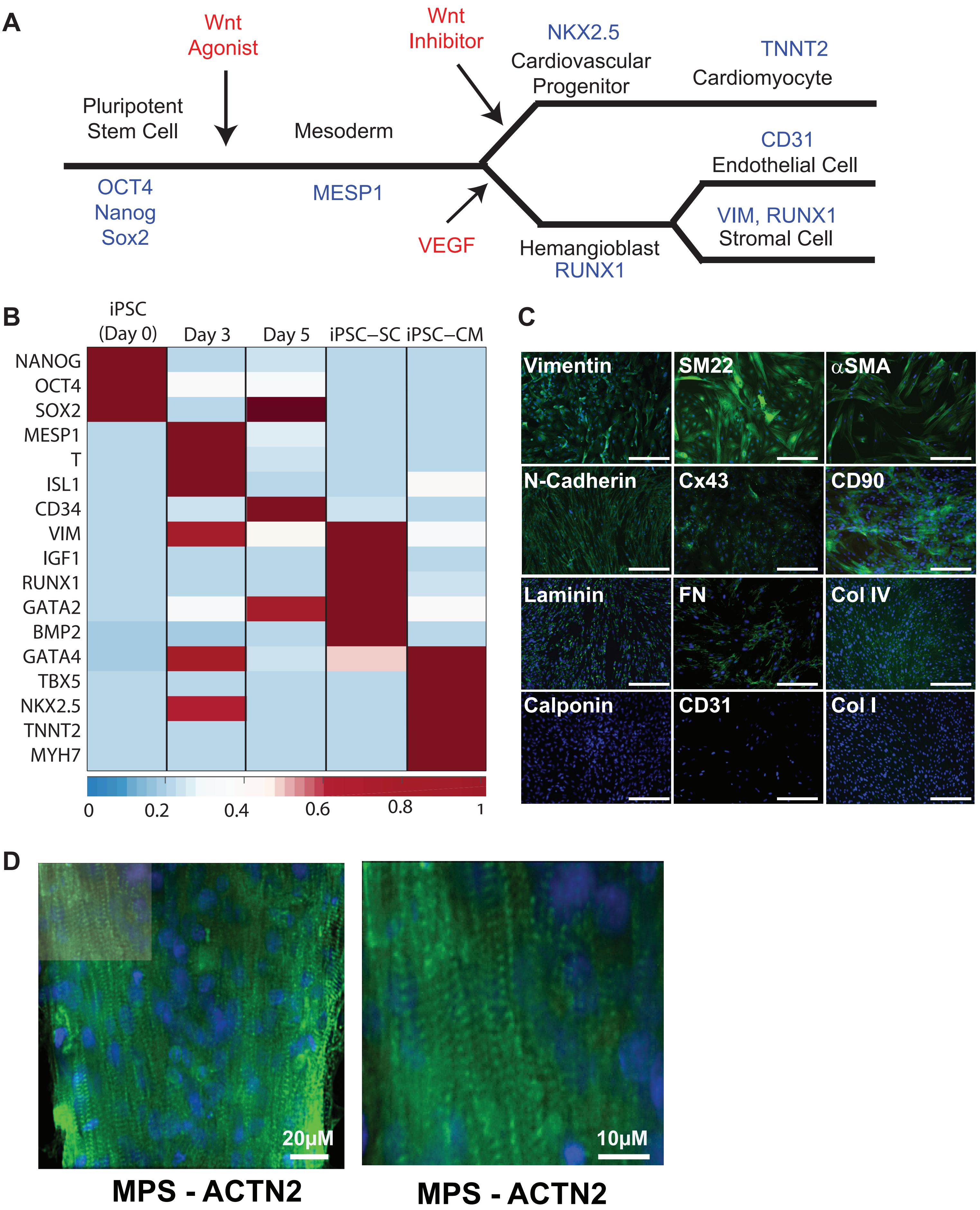
A
Wnt Agonist
Wnt Inhibitor
NKX2.5
Cardiovascular Progenitor
TNNT2
Cardiomyocyte
Pluripotent Stem Cell
Mesoderm
CD31
Endothelial Cell
OCT4
Nanog
Sox2
MESP1
VIM, RUNX1
Stromal Cell
VEGF
Hemangioblast
RUNX1
B
iPSC (Day 0)
Day 3
Day 5
iPSC–SC
iPSC–CM
NANOG
OCT4
SOX2
MESP1
T
ISL1
CD34
VIM
IGF1
RUNX1
GATA2
BMP2
GATA4
TBX5
NKX2.5
TNNT2
MYH7
0
0.2
0.4
0.6
0.8
1
C
Vimentin
SM22
αSMA
N-Cadherin
Cx43
CD90
Laminin
FN
Col IV
Calponin
CD31
Col I
D
20μM
10μM
MPS - ACTN2
MPS - ACTN2

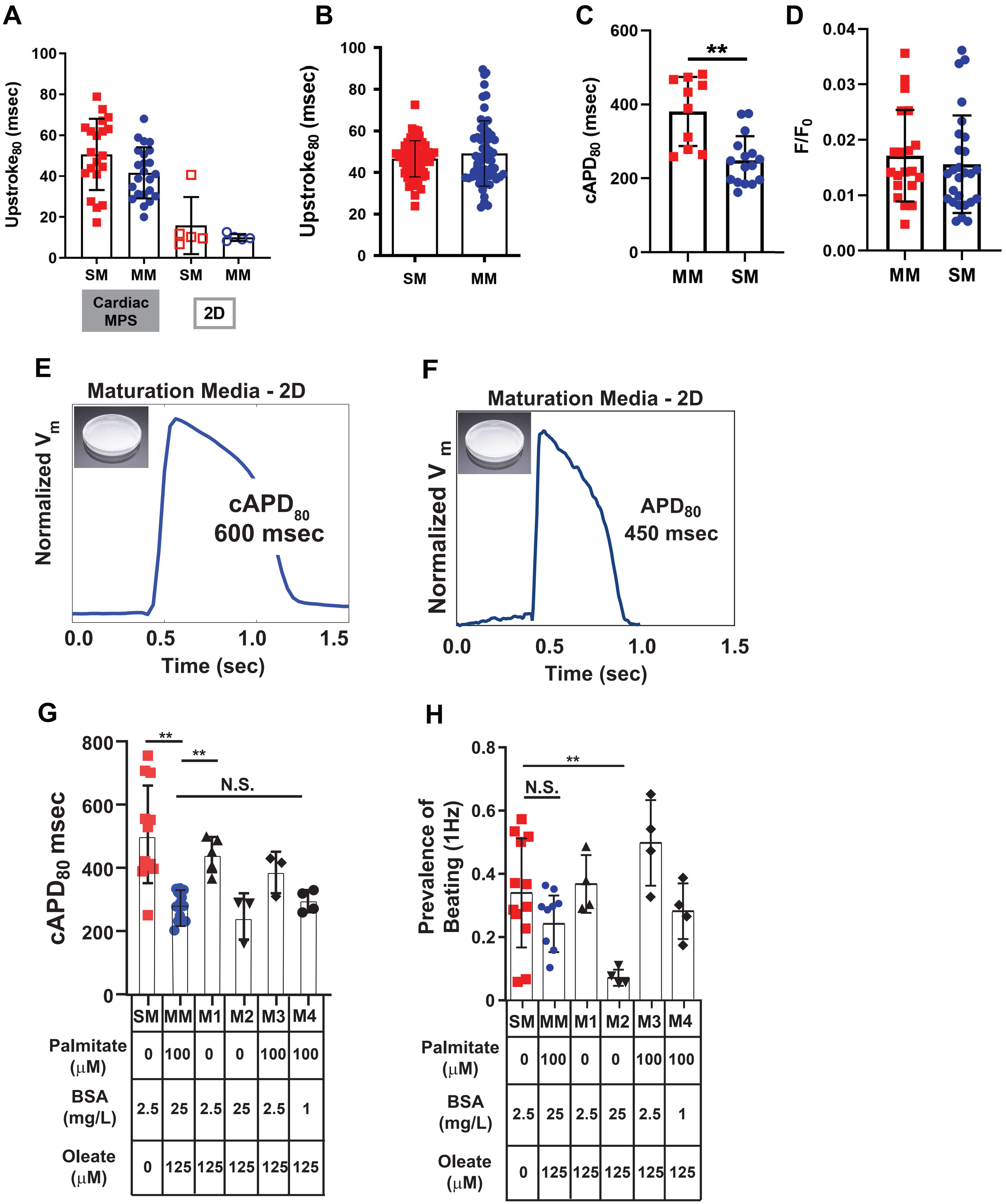


Supp. Figure 3

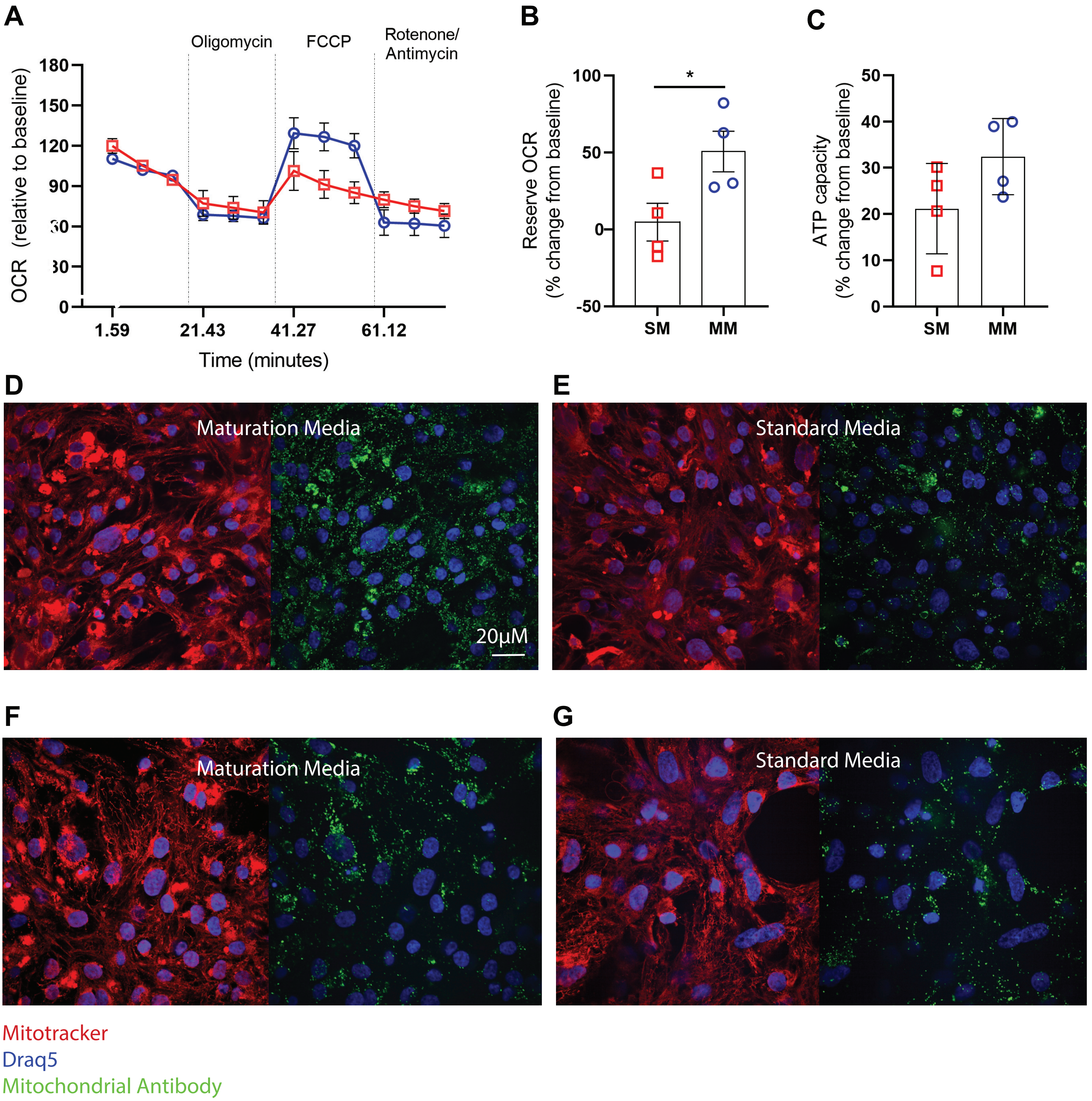


Supp. Figure 4

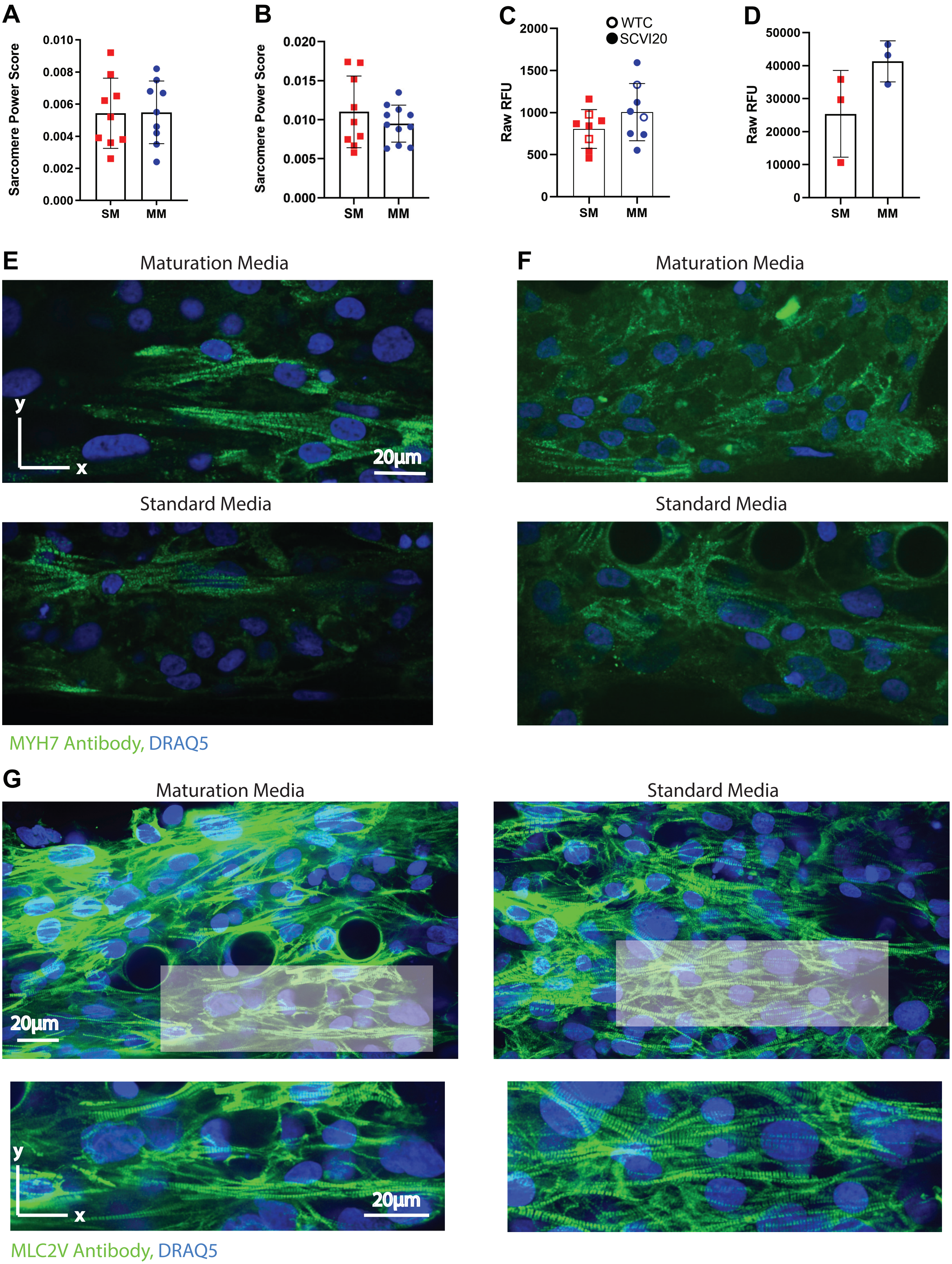


Supp. Figure 5

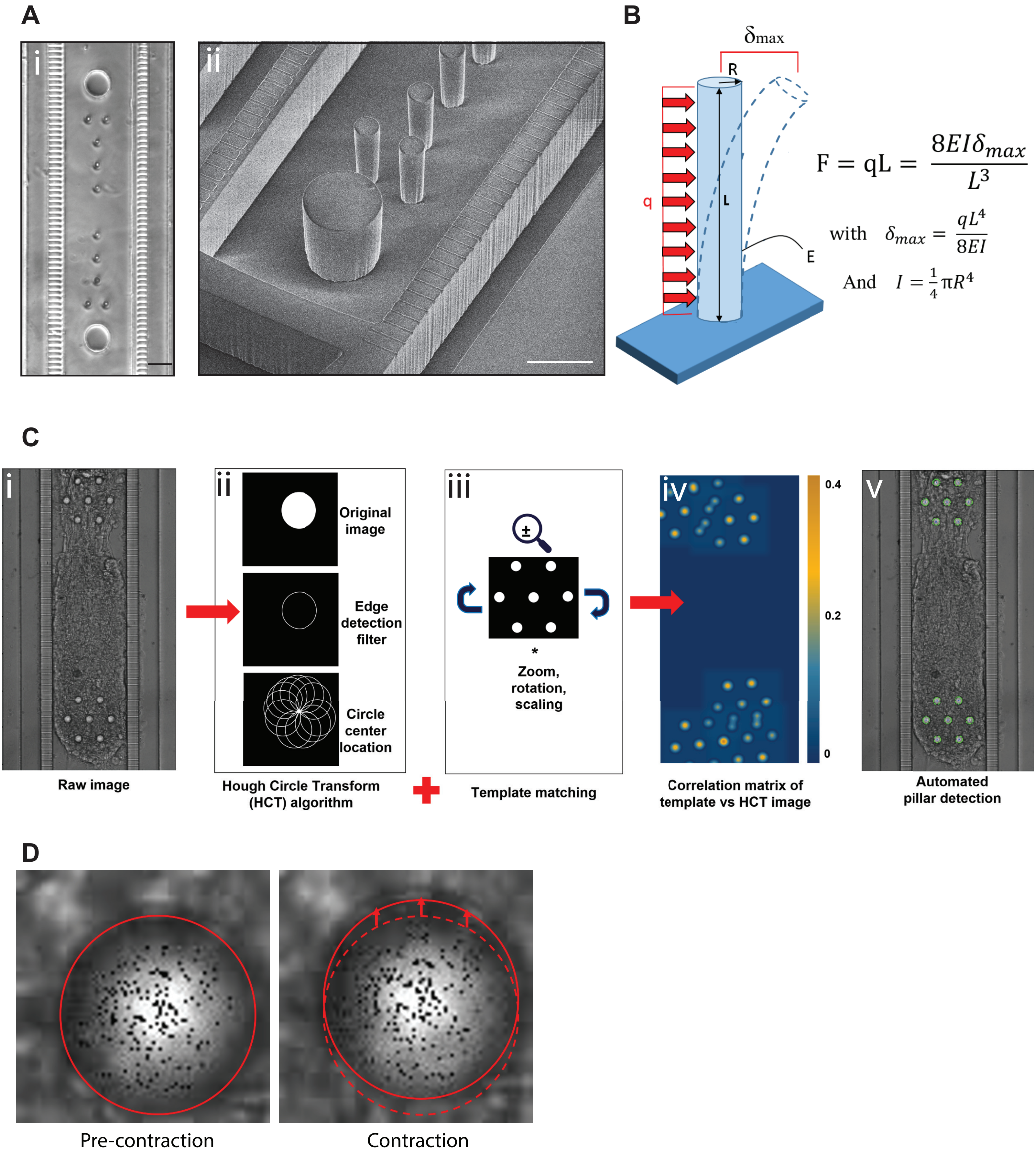


Supp. Figure 6

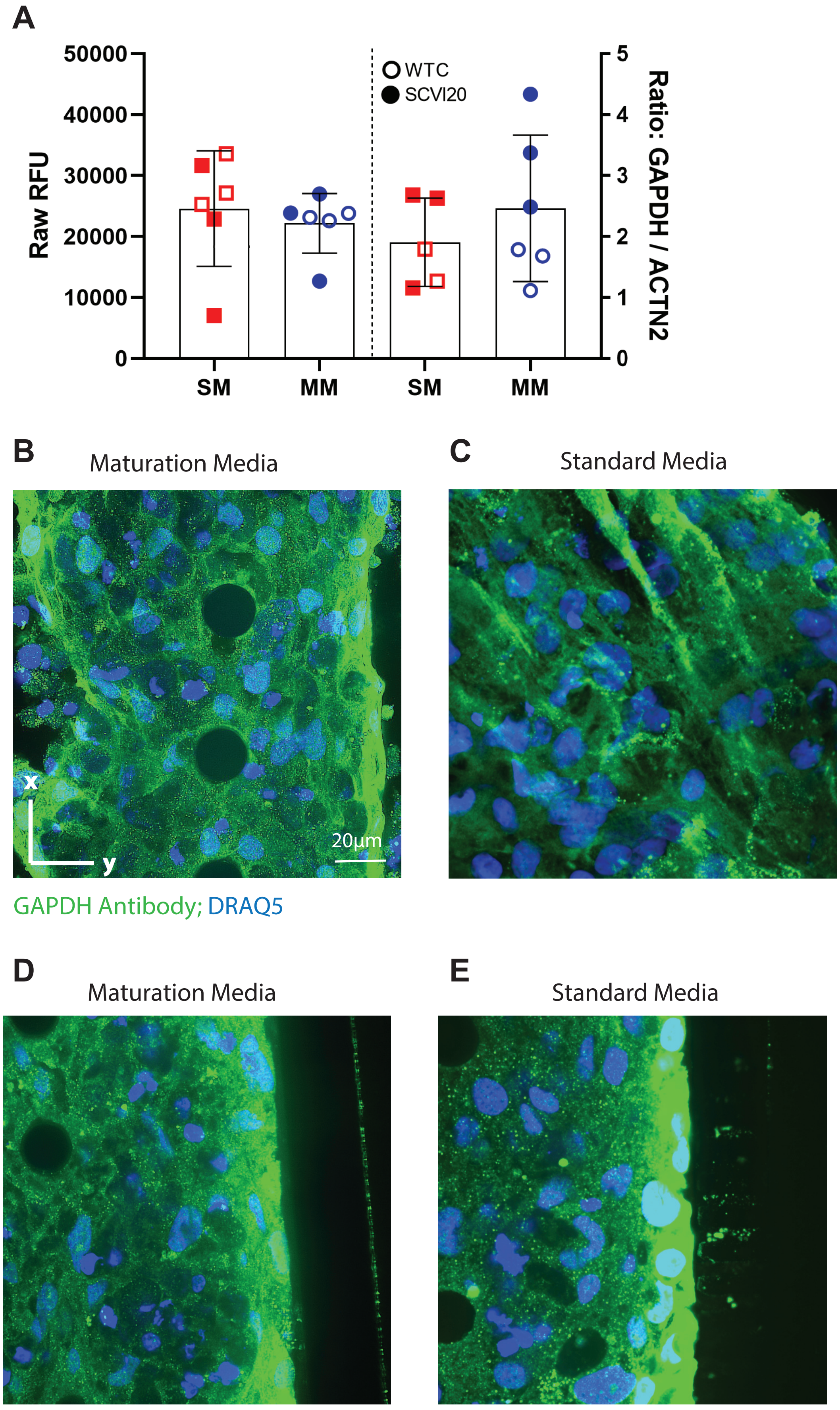


Supp. Figure 7

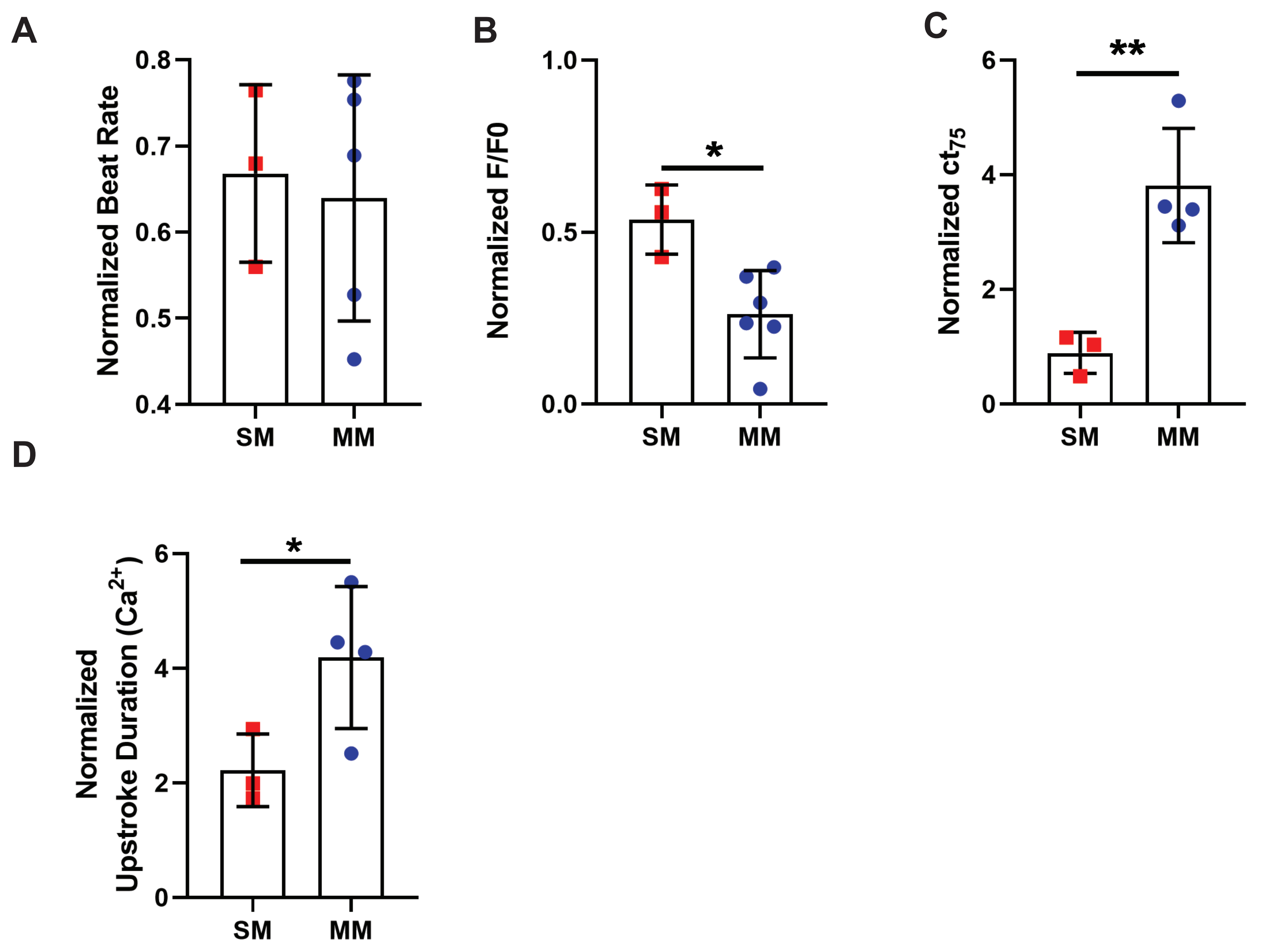


Supp. Figure 8

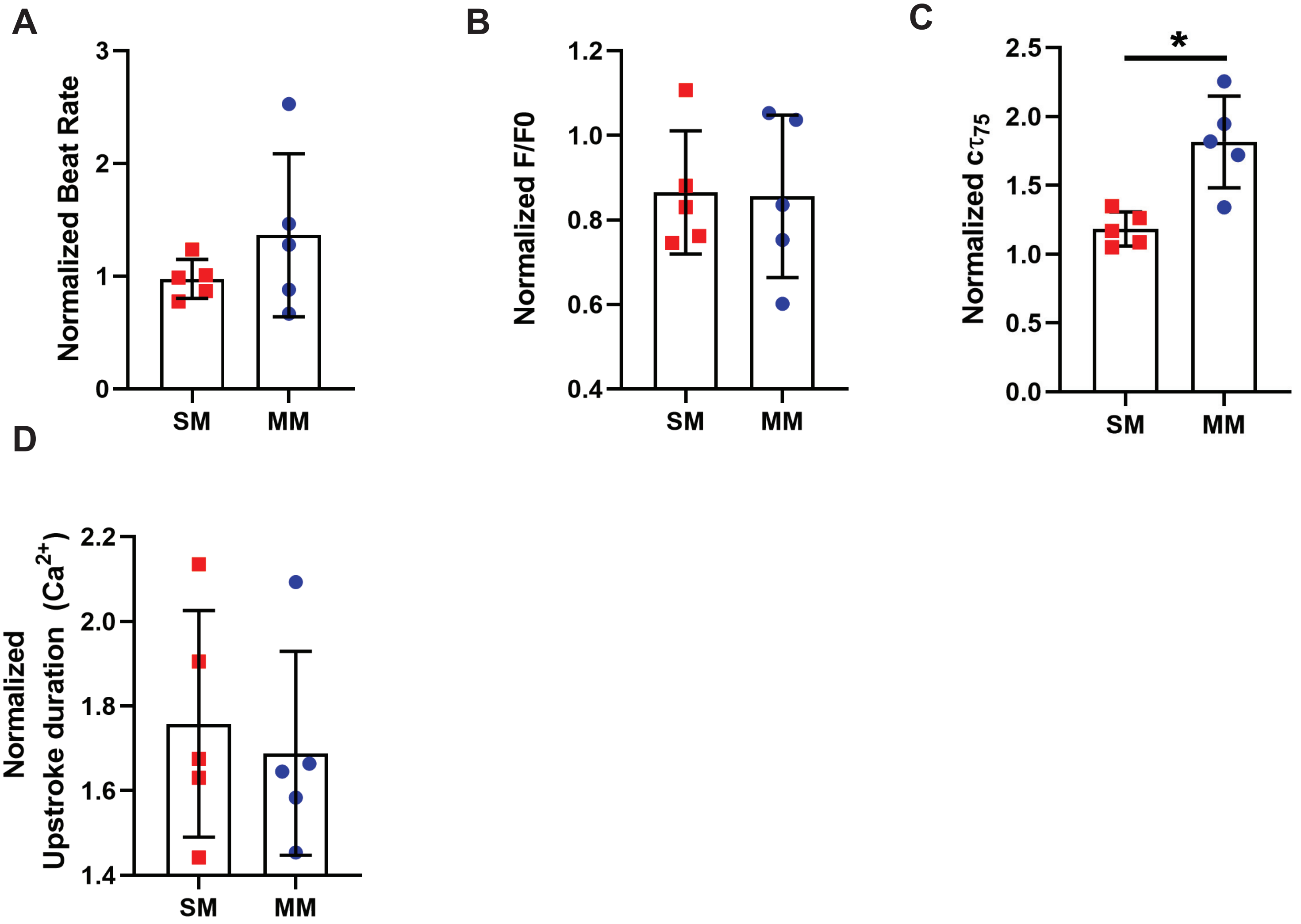


Supp. Figure 9

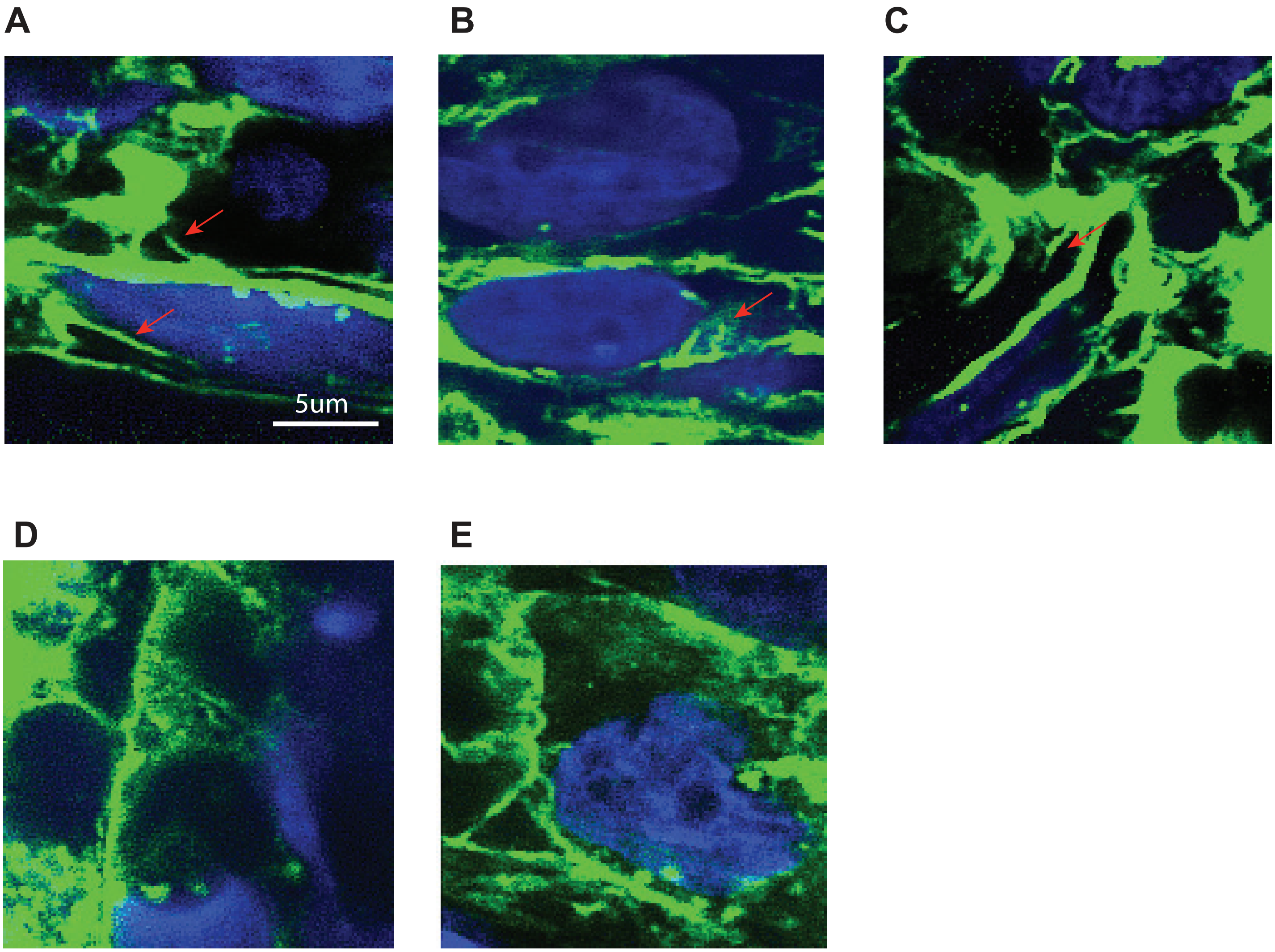


Supp. Figure 10

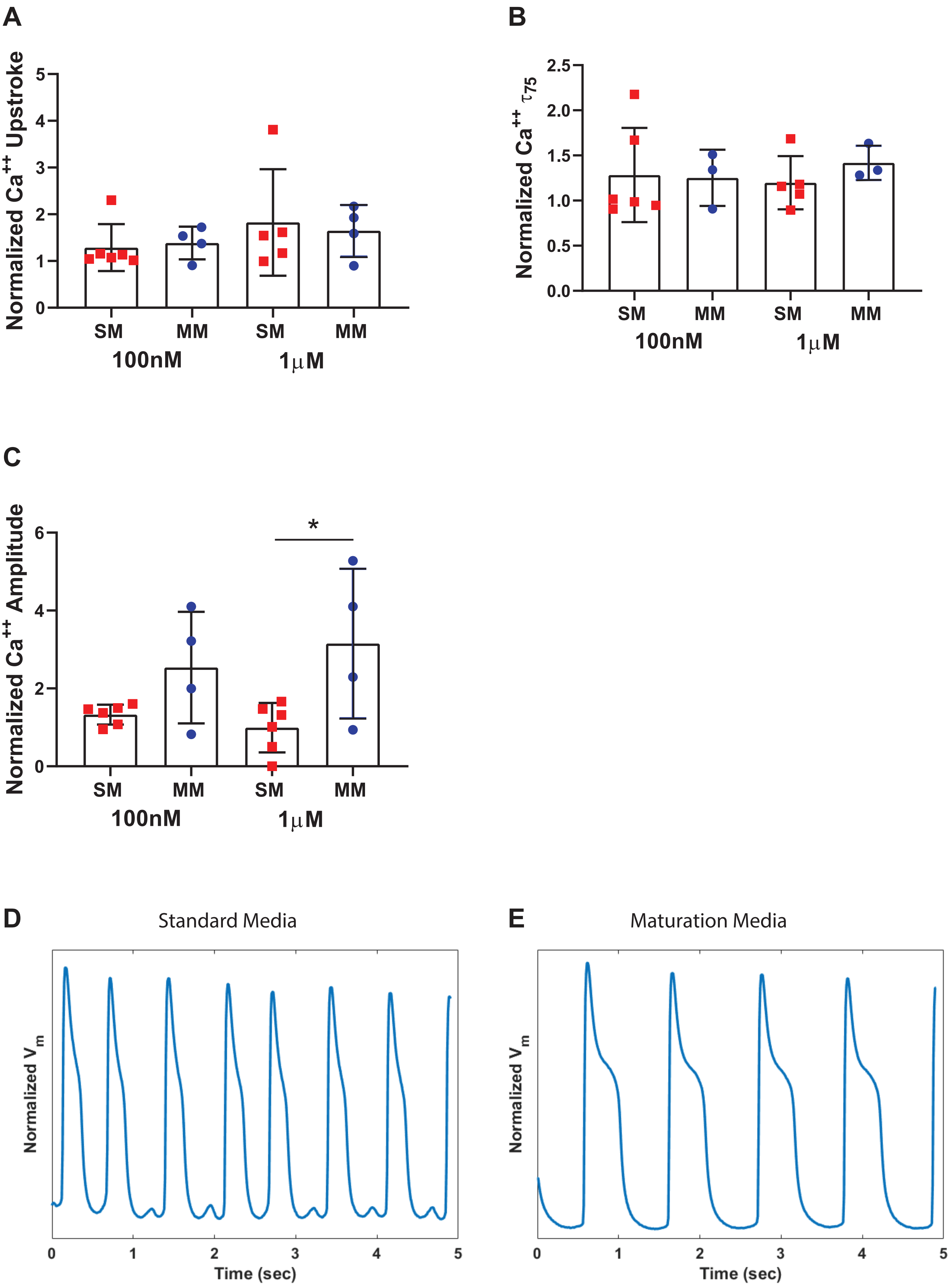


Supp. Figure 11
